# The rice pentatricopeptide repeat protein PPR756 is involved in pollen development by affecting multiple RNA editing in mitochondria

**DOI:** 10.1101/844241

**Authors:** Qiannan Zhang, Yanghong Xu, Jishuai Huang, Kai Zhang, Haijun Xiao, Xiaojian Qin, Linlin Zhu, Yingguo Zhu, Jun Hu

## Abstract

In land plants, the pentatricopeptide repeat (PPR) proteins form a large family involved in post-transcriptional processing of RNA in mitochondria and chloroplasts, which is critical for plant development and evolutionary adaption. Although studies showed a number of PPR proteins generally influence the editing of organellar genes, few of them were characterized in detail in rice. Here, we report a PLS-E subclass PPR protein in rice, PPR756, loss of function of which led to the abolishment of RNA editing events among three mitochondrial genes including *atp6, ccmC*, and *nad7*. Their defective C-to-U transformation then resulted in improper amino acid retention which could cause abortive pollen development. Furthermore, PPR756 could bind to the three target genes directly and interact with three OsMORFs (multiple organellar RNA editing factors): OsMORF1, OsMORF8-1, and OsMORF8-2. The knock-out plants of *PPR756* exhibited retarded growth and greener leaves during the early vegetative stages, along with sterile pollen and lower seed setting at the reproductive stage. These results established a role for PPR756 in rice development, participating in RNA editing of three various transcripts and cooperating with OsMORFs via an editosome manner in rice.

## INTRODUCTION

The pentatricopeptide repeat (PPR) proteins constitute an interesting protein family widely spread in eukaryotes, from algae to humans, although their numbers vary considerably in different organisms (Lurin et al., 2004). This protein family has been suggested to have undergone great expansion when the terrestrial plants came into being andrange from ∼100 (in *Physcomitrella patens*) to over 1000 (in *Selaginella moellendorffii*) (O’Toole et al., 2008;Fujii and Small, 2011). However, other eukaryotic species, such as animals, contain few PPR members, with seven present in humans and six in mice (Lightowlers and Chrzanowska-Lightowlers, 2008;2013). Besides the expansion, a recent study showed that most land plant lineages with high numbers of editing factors have continued to generate novel sequence diversity (Gutmann et al., 2020). Generally, PPR proteins constitute a protein family that is characterized by 2 to about 30 degenerate tandem repeats of about 35 amino acids (Lurin et al., 2004). In general, there are three different types of PPR motifs, including the P motif (35 amino acids), L motif (35 or 36 amino acids), and S motif (31 amino acids), according to which the PPR proteins can be divided into two main classes, P-class and PLS-class. Interestingly, most of the PLS-class PPR proteins contain conserved C-terminal sequences (E, E+ and DYW domain), divided into three subclasses as PLS-E, E+ and DYW subclass, respectively (Small and Peeters, 2000;Lurin et al., 2004). However, based on the alignment of mutiple linear sequence, many distinct PPR motifs definitions were developed, including differing motif borders and numbering schemes (Barkan et al., 2012;Yagi et al., 2013;Yin et al., 2013). Recent study suggested the definition based on the analysis of protein structures could be more accurate (Cheng et al., 2016).

The mitochondria and chloroplasts in plants are semi-autonomous organelles, estimated to contain 2000 and 3000 proteins, but only approximately 40 and 100 genes, respectively, are encoded by these organelles. Massive proteins are encoded by nuclei and imported into these organelles to play important roles in regulation of cellular processes (Mullet, 1988;Sato et al., 1999;Millar et al., 2005). Accumulating research has established that PPR proteins are almost exclusively targeted to the mitochondria or chloroplasts, where they take part in post-transcriptional processing of RNA by altering the RNA sequence, stability, cleavage, splicing, or translation to regulate the organellar gene expression (Dahan and Mireau, 2013;Shikanai and Fujii, 2013).

Different types of PPR proteins have been indicated to perform distinct functions during plant development, such as most P-class PPR proteins that always take part in the progress of organelle transcript cleavage, stability, and translation, while the majority of PLS-class PPR proteins act as editing factors in the post-transcriptional process (Okuda et al., 2007;Saha et al., 2007). For example, PPR10 (Pfalz et al., 2009;Prikryl et al., 2011), ATP4 (Zoschke et al., 2012), CRP1 (Barkan et al., 1994), and PGR3 (Yamazaki et al., 2004) can both stabilize the organelle gene transcripts and activate their translation. Although almost all of the target transcripts are chloroplast ORFs (open reading frame), few are known to affect the mitochondrial gene translation, except for MPPR6, which can influence the choice of start codon and the processing or stabilization of the *rps5* 5’ terminal (Manavski et al., 2012).

It has come to be known that some PPR-type *Rf* genes always belong to the P-class and influence the cleavage of sterility-associated mitochondrial RNAs (Dahan and Mireau, 2013). For instance, in rice, RF1a and RF5 participate in the cleavage of toxic chimeric mitochondrial transcripts contributing to fertility restoration (Wang et al., 2006;Hu et al., 2012). Not only did the P-class PPR proteins take part in the splicing of Group II introns (such as THA8) but several PLS-class proteins were also involved in splicing (such as PpPPR43) (Ichinose et al., 2012;Khrouchtchova et al., 2012). However, in terms of editing factors, the PLS-class proteins may act as the main forces (Kotera et al., 2005). Up to now, a series of PLS-class proteins have been reported to play important roles in the editing of mitochondrial or chloroplast genes. In rice, *OsOGR1, OsSMK1*, and *OsMPR25* have been indicated to participate in the editing of organelle transcripts (Kim et al., 2009;Toda et al., 2012;Li et al., 2014). Previously, we also reported that *OsPGL1* is responsible for the RNA editing of *ndhD* and *ccmFc*, and *OsPPS1* is responsible for *nad3* (Xiao et al., 2018a;Xiao et al., 2018b).

In this study, we characterized a novel PPR gene, *PPR756* (*Os12g19260*), encoding a PLS-E PPR protein, which is associated with the development of rice. PPR756 acted as an editing factor required for the editing of the mitochondrial genes *atp6, ccmC*, and *nad7*, loss of function of which could influence the activities of the mitochondrial electron transport chain complexes and result in dysfunctional pollen.

## MATERIALS AND METHODS

### Plant materials and growth conditions

To construct the RNAi lines, a fragment (ranging from 13 to 354 bp) of *Os12g19260* cDNA was amplified with primers (Table S1) and cloned into the pH7GWIWG(II) vector to construct the RNAi vector. To construct the knock-out lines, we used the CRISPR-Cas9 system and designed two gRNAs (sgRNA1: CGCCCGAAACGAGTACTCCTGG and sgRNA2: GTATCCTGCTACGTGCGGGCTGG) driven by OsU6a and OsU3 for two target sites close to the start codon ATG. The two sgRNA expression cassettes were cloned into Cas9 vector (pYLCRISPR/Cas9P_ubi_-H) for genetic transformation. A series of mutant lines were obtained by two independent transformation experiments. *ppr756-1* and *ppr756-2* were obtained from one transformation, while *ppr756-3* and *ppr756-4* were obtained from the other. The overexpression lines were created by carrying the full length of cDNA of PPR756 without the stop codon fused to the N-terminus of tags, driven by the ubiquitin promoter in the pCAMBIA1301 backbone. All the constructs were transformed into the calli of *Oryza sativa* L. *japonica* Zhonghua 11 (ZH11) mediated by *Agrobacterium tumefaciens*. All the transgenic plants were grown in Hubei province and Hainan province, China, and in a growth chamber in Wuhan, Hubei province, under proper management.

### Subcellular localization of PPR756

For transient expression, the full-length coding sequence of PPR756 was cloned into HBT-sGFP vector under the control of a CaMV 35S promoter. The protoplasts were extracted from 1-week-old etiolated rice seedlings and then transformed with 10–20 μg plasmids according to the procedure described in (Xiao et al., 2018b). Mito Tracker Red (Invitrogen) was used as a mitochondrial specific dye. The organelle and GFP signals were detected with a Leica microscope (DM4000 B, Germany) using different excitation wavelengths. In addition, Western blotting was performed to confirm the subcellular localization of PPR756, in which the mitochondria and chloroplasts were extracted from the PPR756 OE (overexpression) lines. Then the signals were detected by antibodies, including Flag (DIA-AN, Wuhan, China), VDAC, RbcL (Agrisera, Vannas, Sweden), actin, and histone (ABclonal). Moreover, in order to enhance the Flag signals in the total protein, different amounts of proteins were loaded as Total: Mitochondria: Chloroplasts=5:1:1.

### RNA Extraction, reverse transcription PCR and Real-time PCR

The total RNA was extracted using Trizol reagent (Invitrogen) according to the manufacturer’s instructions. The RNA was resuspended in RNase-free water and treated with RNase-free DNase I (Thermo Fisher). After digestion, PCR was performed to confirm the elimination of DNA contamination. Approximately 5 μg of the digested total RNA was reverse-transcribed using the M-MLV reverse transcriptase (Invitrogen) and random primers (Thermo Fisher) to obtain the first-strand cDNA. The real-time PCR was performed on a Lightcycler 480 (Roche) using the SYBR Green I Master PCR kit with gene-specific primers. Ubiquitin and actin served as controls for gene expression.

### RNA editing analysis

The primers for PCR and sequencing (Table S1) were designed according to the reported editing genes in mitochondria and chloroplasts. Then the cDNAs obtained from reverse-transcription of the WT and transgenic lines were used as the template for PCR with different primers combinations, and the PCR products were sequenced. For the editing analysis, the sequencing results of RNA editing of transgenic lines were compared with those of the WT to detect the change of RNA editings. Three repeated experiments were performed in different independent lines with biological duplications. The synthesis of primers and sequencing were accomplished in TSINGKE Biological Technology.

### Histochemical analysis of GUS activity

To further detect the temporal and spatial expression pattern of *PPR756*, a 1963 bp fragment including the promoter cassette of *PPR756* was amplified from ZH11 and cloned into the vector pCAMBIA1391 in order to drive the GUS reporter gene expression. The vector was transformed into the calli of ZH11 to obtain the transgenic plants. Then, the GUS activity of various tissues in the transgenic plants was measured according to the methods of our previous study (Xiao et al., 2018b).

### Yeast two-hybrid assay

The full-length cDNA of *PPR756* was amplified and cloned into the bait vector pGBKT7, while *OsMORF1, OsMORF2, OsWSP1, OsMORF3, OsMORF8-1, OsMORF8-2*, and *OsMORF9* were cloned respectively into the prey vector pGADT7. These constructs were then co-transformed into the yeast strain AH109 in corresponding pairs as in a previously described method (Hu et al., 2012).

### Bimolecular fluorescence complementation assays

For the bimolecular fluorescence complementation (BiFC) analysis, the full-length cDNA of *PPR756* was fused into the C-terminus fragment of yellow fluorescent protein (YFP) in the pUC-SPYCE vector, and OsMORF proteins were fused into the N-terminus fragment of yellow fluorescent protein (YFP) in the pUC-SPYNE vector. Constructs were co-transformed into rice protoplasts in pairs, and the signals were observed using a Leica microscope (DM4000 B, Germany) in bright and fluorescent fields as described previously (Hu et al., 2012).

### Determination of chlorophyll content

Acetone extraction technology was used to isolate chlorophyll. The extracting solution was absolute ethyl alcohol/acetone/water at a ratio of 4.5:4.5:1. Fresh leaves (0.5 g) derived from the wild-type (WT), over-expression (OE) line, and knock-out (KO) line were cut into 1 cm slices and put into 10 mL of extracting solution in total darkness for approximately 20 h until the leave slices became white. A 200 μl sample of the pigment solution was taken to measure the absorption values, with the extracting solution as a control, using the Tecan Infinite M200 (Switzerland), and the chlorophyll contents were calculated according to the formula described previously (Arnon, 1949).

### Recombinant protein expression and RNA electrophoresis mobility shift assays

The recombinant protein was created with a fusion of MBP in the PPR756^41-756^ N-terminus and 6xHis in the C-terminus, and was purified across two columns equipped with Ni^2+^ affinity resin (Ni-NTA Resin, GenScript) and MBP (PurKine MBP-Tag Dextrin Resin 6FF, Abbkine) in turn. The control fusion protein containing only MBP and His tags was purified as well. RNA probes (Table S1) were synthesized and labeled with 6-FAM at the 5’ end by GenScript (Nanjing, China). For the RNA electrophoresis mobility shift assays (REMSAs), the dialyzed recombinant protein was incubated with 100 fmol RNA probes in a 10× binding buffer (100 mM HEPES PH = 7.3, 200 mM KCl, 10 mM MgCl_2_, 10 mM DTT) condition and reacted in a 20 μl system for 30 min at 25 °C with 10 units of RNasin, followed by separation on 8% native acrylamide gels in a 0.5× TBE buffer. After electrophoresis, the gels were screened using a Typhoon Trio Imager (GE).

### RNA Immunoprecipitation-PCR (RIP-PCR)

RIP is a well-developed technology to detect protein–RNA interactions in vivo and involves the immunoprecipitation (IP) of a target protein followed by purification of the associated RNA. Here, the cell lysate was obtained from 2-week-old OE line seedlings, which were cross-linked in 0.1% formaldehyde in advance to strengthen protein–RNA interactions. The seedlings were ground in liquid nitrogen, and the powder was resuspended in extraction buffer (Tu et al., 2015). Then, we used the anti-Flag antibody to immunoprecipitate the PPR756-Flag fusion protein from the supernatant with IgG as a negative control. After immunoprecipitation, the associated RNA was extracted using Trizol and reverse-transcribed with random primers to obtain cDNA. The cDNA was further detected by quantitative PCR (qPCR).

### Scanning electron microscopy analysis

The samples for SEM analysis were prepared according to a previous study (Xiao et al., 2018a). The seedlings were cut in 1 cm long sections, and mature spikelets were immersed in 2.5% glutaraldehyde buffer for 18 h at 4 °C as immediately after separation from the plants. The samples were dehydrated using different concentrations of ethyl alcohol at room temperature every hour, and then the dehydrating agent was replaced in order to dry the samples at the critical point. Lastly, the samples were sputter coated with gold in an ion sputter (E-100, Japan) and observed with a scanning electron microscope (Hitachi S-3000N, Japan).

### Detection of mitochondrial complex activity

The activities of mitochondrial electron transport chain complexes, complex I, III, IV and V were measured using the electron transport chain complex assay kits from Solarbio Life Science according to the instructions (Beijing Solarbio Science & Technology).

## RESULTS

### PPR756 is essential for rice development

In previous reports, to identify the function of PPRs in rice, we generated a series of RNAi transgenic lines of several PPR genes with the background of ZhongHua11 (ZH11, *Oryza sativa, L. japonica*) and characterized OsPGL1 and OsPPS1 (Xiao et al., 2018a;Xiao et al., 2018b). In this study, we investigated novel independent RNAi lines, in which the expression of *Os12g19260* was knocked down solidly (Supplementary Figure 1C). Since *Os12g19260* encodes a PPR protein with 756 amino acids, we identified it as *PPR756* in this study. These lines displayed pleotropic phenotypes compared with the WT, including delayed development, more green leaves, and smaller leaf angles in the early vegetative stage (Supplementary Figures 1A–E).

Based on these interesting phenotypes, we applied the CRISPR-Cas9 system to create knock-out (KO) mutant lines using the background of ZH11 to confirm the function of PPR756. We totally obtained several independent mutant lines, which exhibited same phenotypes (Supplementary Figure 5A-F), two lines displayed premature translation termination were chosen and named *ppr756-1* and *ppr756-2* (Figure 1A, Supplementary Figure 2A), rest of them were named *ppr756-3* and *ppr756-4*. In order to explore the function of *PPR756* in detail, we also generated over-expression (OE) transgenic lines, in which the CDS of PPR756 was driven by the ubiquitin promoter and fused with the Flag and cMyc tags (Supplementary Figure 2B). We checked the expression of PPR756 in three independent OE lines, named PPR756-OE-1 to PPR756-OE-3. The relative expression of PPR756 was elevated in all OE lines, especially in PPR756-OE-2 (Supplementary Figure 2C). Assessment of protein accumulation was conducted simultaneously. Western blotting confirmed that PPR756 was successfully expressed in the transgenic line (Supplementary Figure 2D). Based on the above detection and confirmation, we selected PPR756-OE-2 and *ppr756-1* for further analysis in this study.

**FIGURE 1.**
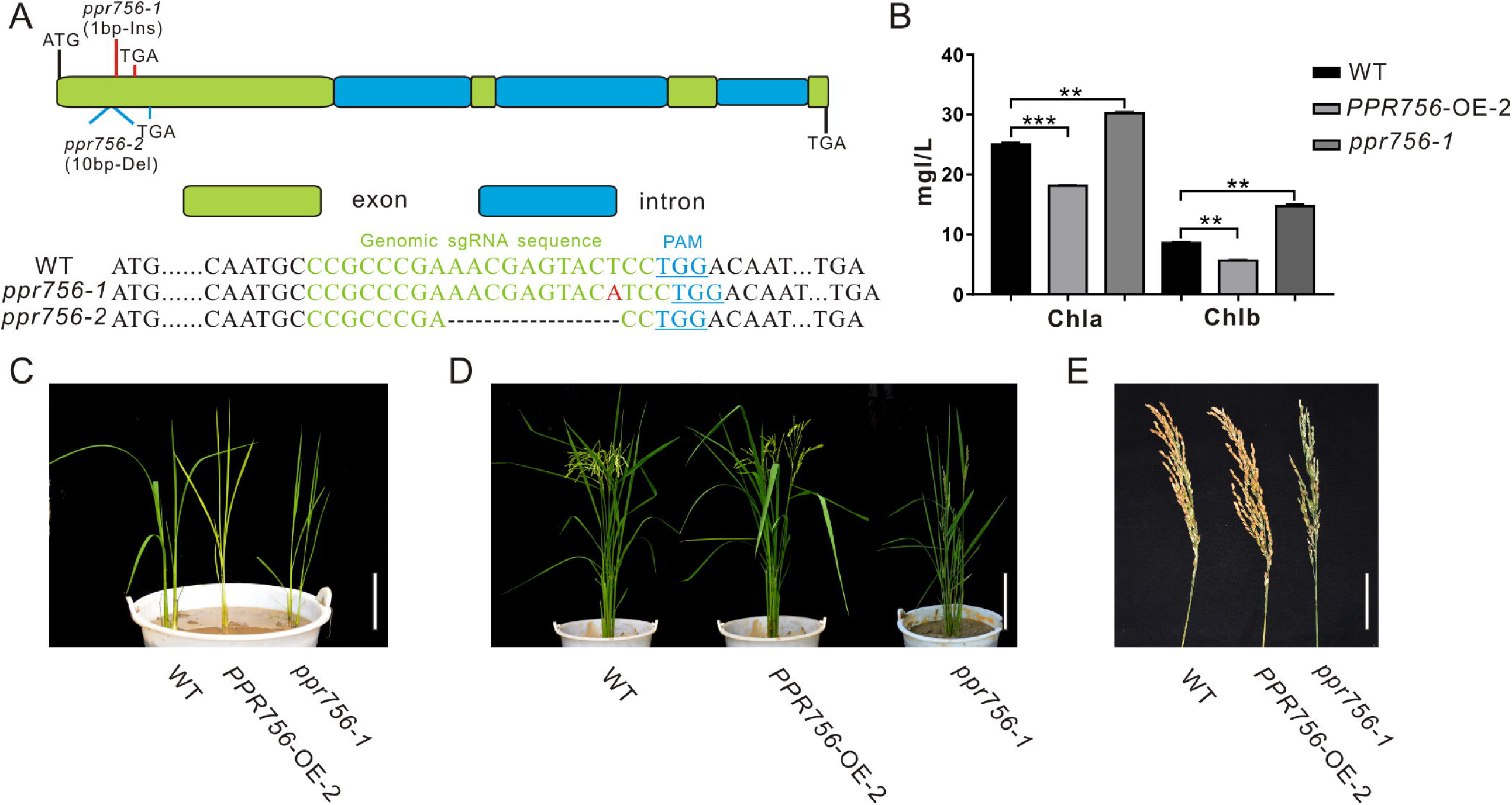
Phenotypic characterization of the *PPR756* lines. **(A)** The structure of *PPR756* and *ppr756* mutant. Two target sites were chosen, and the obtained *ppr756* mutant lines showed premature translation termination with 1 bp insertion in *ppr756-1* line and 10 bp deletion in *ppr756-2* line. **(B)** The different chlorophyll contents of leaves in the wild-type (WT), the *PPR756* OE line, and *ppr756* mutant. The bars represent the SD of three independent biological replicates, and the unit of content is mg/L. **(C–D)** Phenotypic comparison of the seedlings and mature plants of the wild-type (WT), the *PPR756* OE line, and *ppr756* mutant. Bars, 5 cm for **(C)** and 20 cm for **(D)**, respectively. **(E)** Comparison of the panicles in wild-type (WT), the *PPR756* OE line and *ppr756* mutant. Bars, 5 cm. Quantitative data were means ± SD based on three independent experiments (Student’s t-test; *, *P* < 0.05; **, *P* < 0.01; ***, *P* < 0.001).

Since the leaf color showed obvious visible phenotypic differentiation, we first measured the chlorophyll contents of PPR756-OE-2 and *ppr756-1*. Data showed chlorophyll a (Chla) and chlorophyll b (Chlb) were highly accumulated in KO mutants and reduced in OE lines compared with the WT, which explained the change of leaf color (Figure 1B, C), but the mechanism causing the phenotype remained to be explored. Loss of function of PPR756 also resulted in erect leaves and lower seed setting, while the OE line presented the opposite phenotypes, especially with a larger panicle compared to the WT (Figure 1D, E). All of these results indicated that PPR756 has important roles in rice development and affects the yield in rice.

### PPR756 especially influences pollen fertility

To make it clear whether the lower seed setting resulted from male sterility or female sterility, we first tested the pollen fertility by I_2_-KI staining. The pollen became sterile when PPR756 was null (Figure 2A). To explore the underlying mechanism, we detected the morphological details of the anthers and pollen by SEM. The observations of the anthers showed a curly anther base in the KO mutant line compared with the WT and OE line (Figure 2B). The observation of pollen revealed that most of the pollen grains in the KO mutant were distorted and shrunken, while those in the WT and OE lines were spherical and regular (Figure 2C, D). All of these data indicated that the dysfunctional development of pollen in the KO mutant accounted for the reduced seed setting rate, implying that PPR756 is required for pollen development in rice.

**FIGURE 2.**
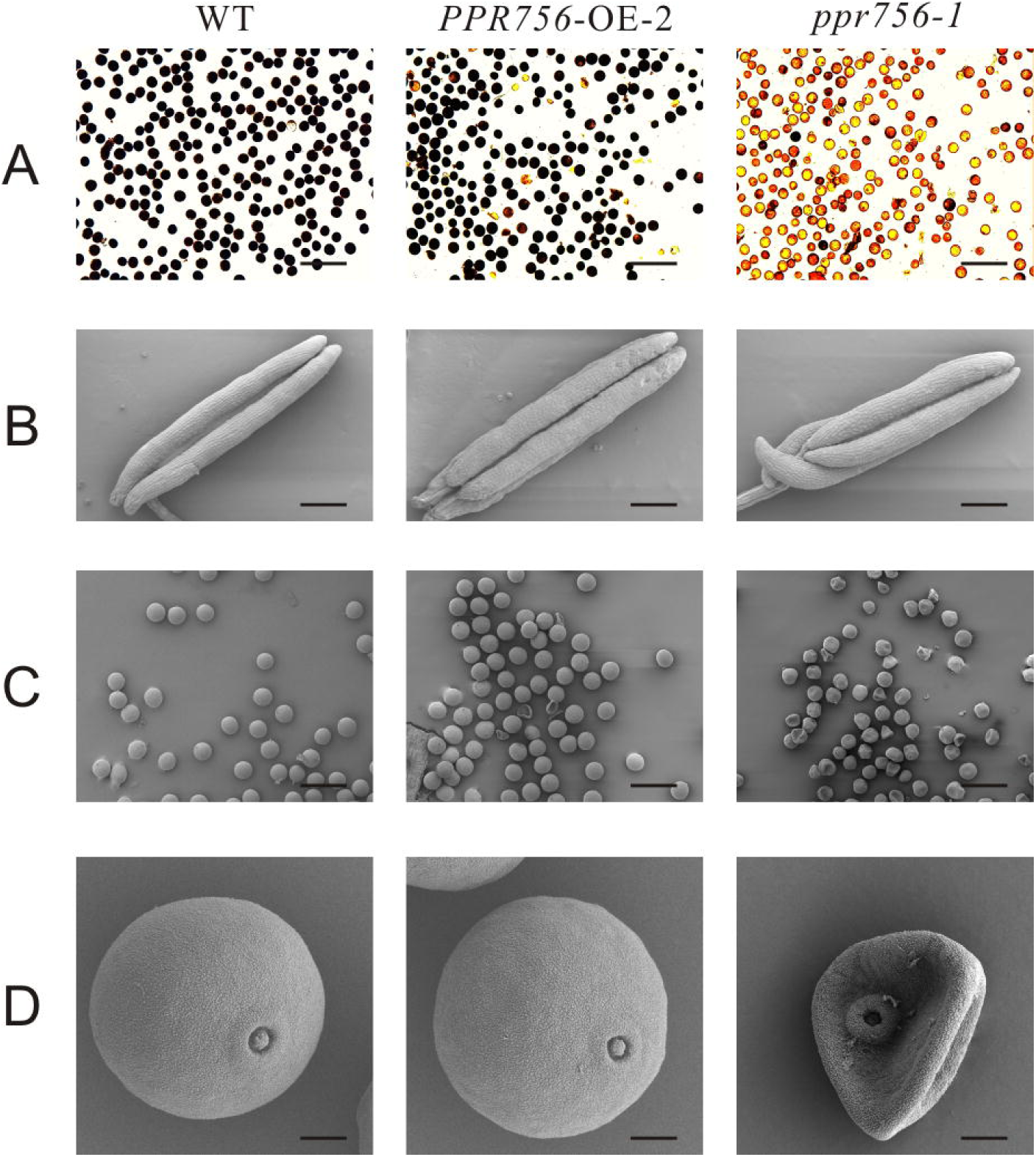
Fertility detection and scanning electron microscopy observations. **(A)** Comparison of pollen fertility of WT, the *PPR756* OE line, and *ppr756* mutant by potassium iodide (1% I_2_-KI) staining. The darkly stained pollen is fertile, whereas the lightly or non-stained pollen is sterile. Bars, 200 μm. Representative results from three independent experiments. **(B)** Scanning electron microscopy (SEM) observation of anthers of WT, the *PPR756* OE line, and *ppr756* mutant. Bars, 200 μm. Representative results from three independent experiments. **(C, D)** Scanning electron microscopy (SEM) observation of pollens of the *PPR756* OE line and *ppr756* mutant. Bars, 100 μm for **(C)** and 10 μm for **(D)**. Representative results from three independent experiments.

### PPR756 encodes a PPR-E subclass protein

Mostly, PPR proteins are translated without introns (O’Toole et al., 2008), though PPR756 consists of four exons and three introns based on the sequence analysis (Figure 1A). PPR756 was predicted to contain 19 putative PPR motifs by Basic Local Alignment Search Tool (BLAST) with the previously reported variant reference sequences (Cheng et al., 2016). However, PPR756 is a unique member of PLS-class, containing 13 successive SS motifs followed by P1-L1-S1-P2-L2-S2-E1-E2 and is considered as a PLS-E subfamily protein (Supplementary Figure 3A). Furthermore, these PPR motifs in PPR756 showed high similarity especially on the functional sites (Supplementary Figure 3B).

To further explore the evolution of PPR756 in plants, a phylogenetic tree was constructed based on the protein sequence alignment from 22 other species, which exhibited over 55% similarity. All of these homologous plants were vascular, with most being monocots and only one dicot (Supplementary Figure 3C). Moreover, the homologous plants in the top five species were further analyzed, with results showing extreme conservation in these monocots (Supplementary Figure 3D). Therefore, data suggested PPR756 may play a conservative role in different plant species, especially in monocots.

### Subcellular localization of PPR756

The majority of PPR proteins are predicted to localize either in mitochondria or chloroplasts (Lurin et al., 2004). Up to now, the characterized PPR protein also confirmed this view, and some of them were dual-localized to both mitochondria and chloroplasts. Bioinformatic analysis of the PPR756 protein sequence using the TargetP 1.1 server (http://www.cbs.dtu.dk/serivices/TargetP/) indicated that PPR756 is dual-localized to chloroplasts and mitochondria. To determine the actual subcellular localization of PPR756 in vivo, we applied the p35S:PPR756-sGFP construct encompassing the full-length cDNA of PPR756 lacking the stop codon fused with GFP (green fluorescent protein) driven by the constitutive CaMV 35S promoter. Then, the recombinant vector and the empty p35S:sGFP vector (control) were transiently expressed in the rice protoplasts. The GFP fluorescent signals were detected by confocal microscopy, showing that the GFP signals were well overlapped with the indicator signal of mitochondria (Figure 3A).

**FIGURE 3.**
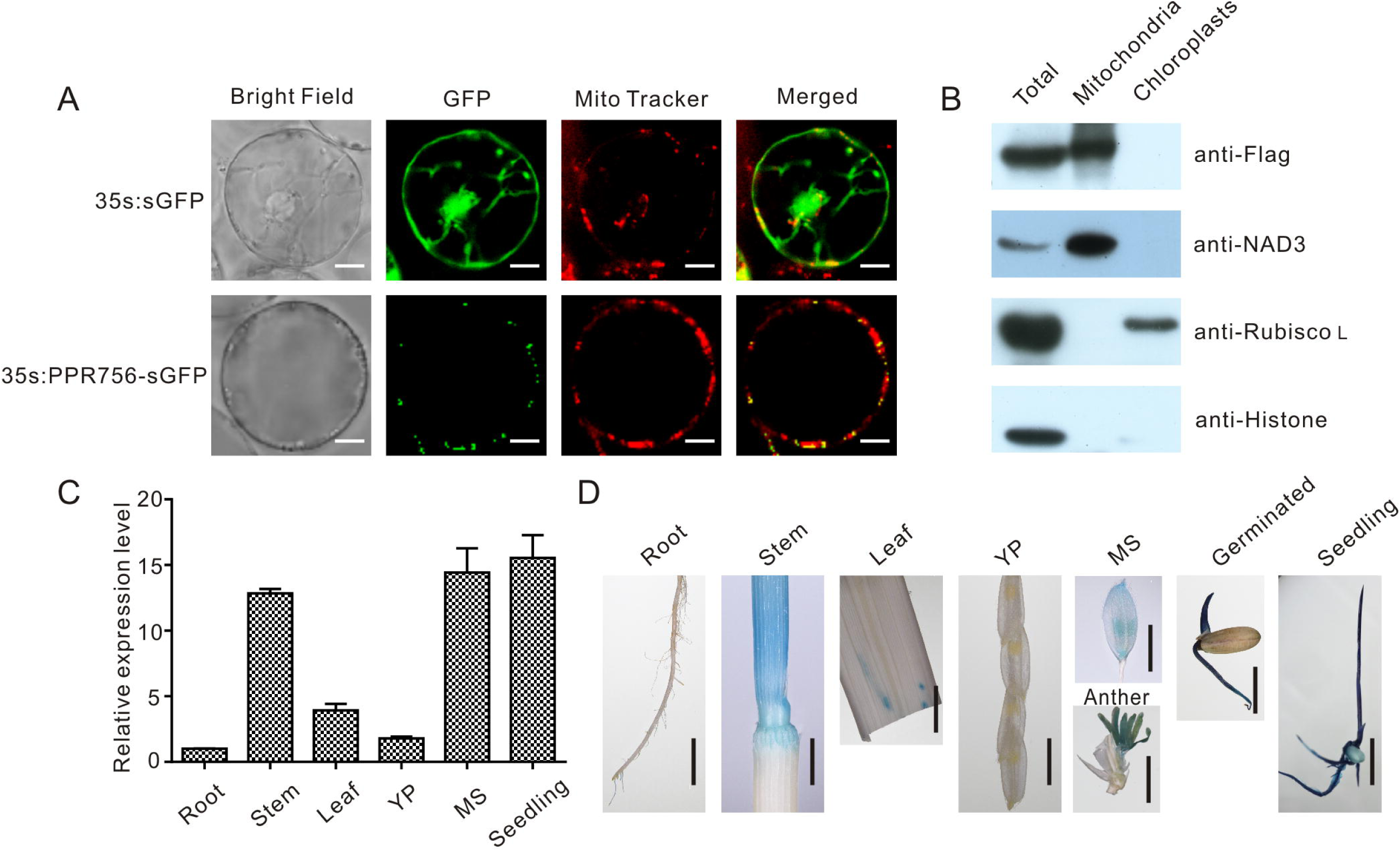
Subcellular localization and expression pattern of PPR756. **(A)** The vectors 35s:sGFP and 35s:PPR756-sGFP were transferred into the rice protoplasts for transient expression. The green fluorescent protein (GFP) was detected and merged with the signal of MitoTracker, a mitochondria indicator. Bars, 5 μm. **(B)** Total proteins, mitochondrial proteins, and chloroplast proteins of transgenic plants were isolated for immunoblot assays to evaluate the subcellular localization of PPR756. Antibodies against NAD3, Rubisco L, and histone were used as indicators for the mitochondria, chloroplasts, and nucleus, respectively. The Flag signal represents the fusion protein PPR756-Flag. **(C)** Quantitative real-time PCR analysis of *PPR756* in various WT tissues. Rice *Actin* gene was used as a control. Quantitative data were means ± SD based on three independent experiments. **(D)** Histochemical staining analysis of *PPR756* promoter–GUS reporter gene in various tissues. YP, young panicle; MS, mature spikelet; Bars, 0.5 cm.

Moreover, the Flag tag was used as a marker in the OE transgenic lines to observe the subcellular location of PPR756. An immunoblot assay with an anti-Flag antibody was then performed to detect the PPR756-Flag fusion protein in the chloroplasts and mitochondria extracted from the OE lines. An obvious signal detected in the mitochondria extract confirmed that PPR756 was mainly targeted to mitochondria, while signal was hardly observed in the chloroplast (Figure 3B). However, PPR756 can interact with OsMORF8 which localizes in both mitochondria and chloroplast (Figure 7A and B), we did not exclude the possibility of PPR756’s location in chloroplast.

**FIGURE 4.**
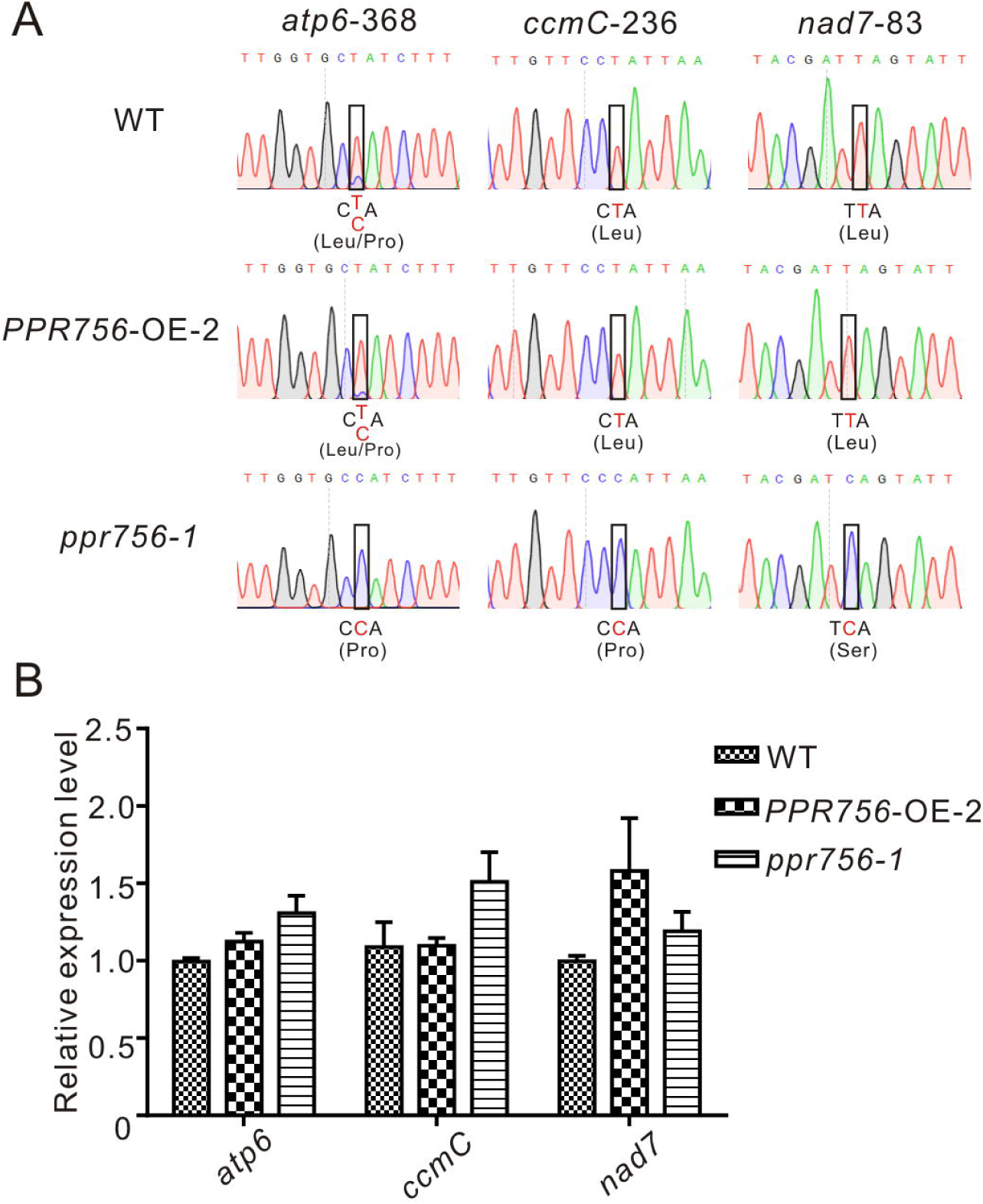
PPR756 is indispensable for mitochondrial gene editing. **(A)** RNA editing analysis of *atp6-368, ccmC-236, nad7-83* in the WT, *PPR756* OE line, and *ppr756* mutant. **(B)** Transcript level determination of the three target genes in the WT, *PPR756* OE line, and *ppr756* mutant. Data are means from three independent biological replicates.

**FIGURE 5.**
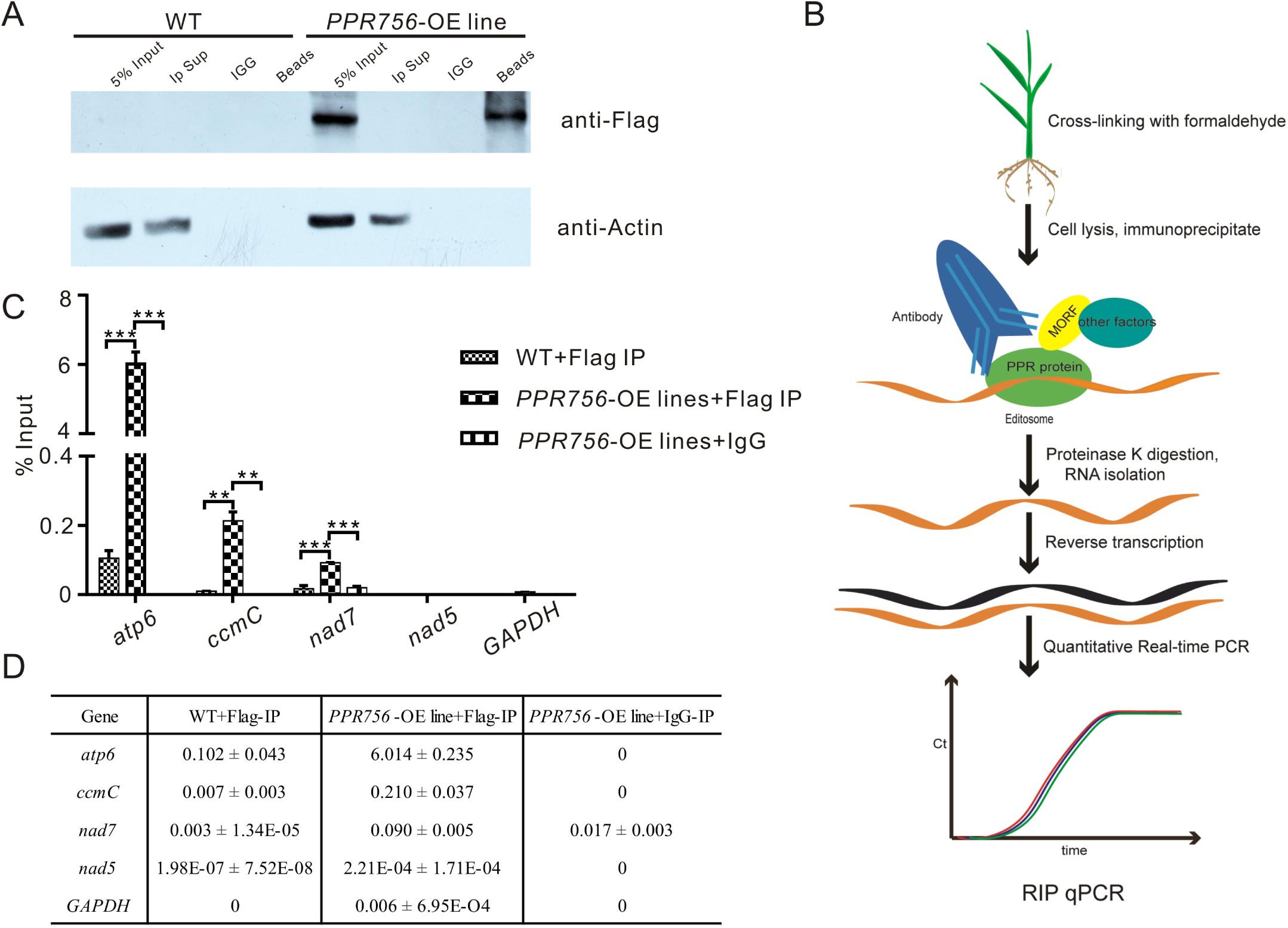
PPR756 can directly bind to its target genes *in vitro* and *in vivo*. **(A)** Western blots were carried out to validate the IP efficiency. Actin was used as a control, and the IgG served as negative control. **(B**) The flowchart of RIP-qPCR experiment. **(C)** Quantitative real-time PCR of *atp6, ccmC*, and *nad7*. The *nad5* and *GAPDH* were used as control genes. Bars represent the SDs from three independent biological replicates. **(D)** The percentage binding rate of three target genes and two control genes in the RIP-qPCR according to the input. Quantitative data were means ± SD based on three independent experiments (Student’s t-test; *, *P* < 0.05; **, *P* < 0.01; ***, *P* < 0.001).

**FIGURE 6.**
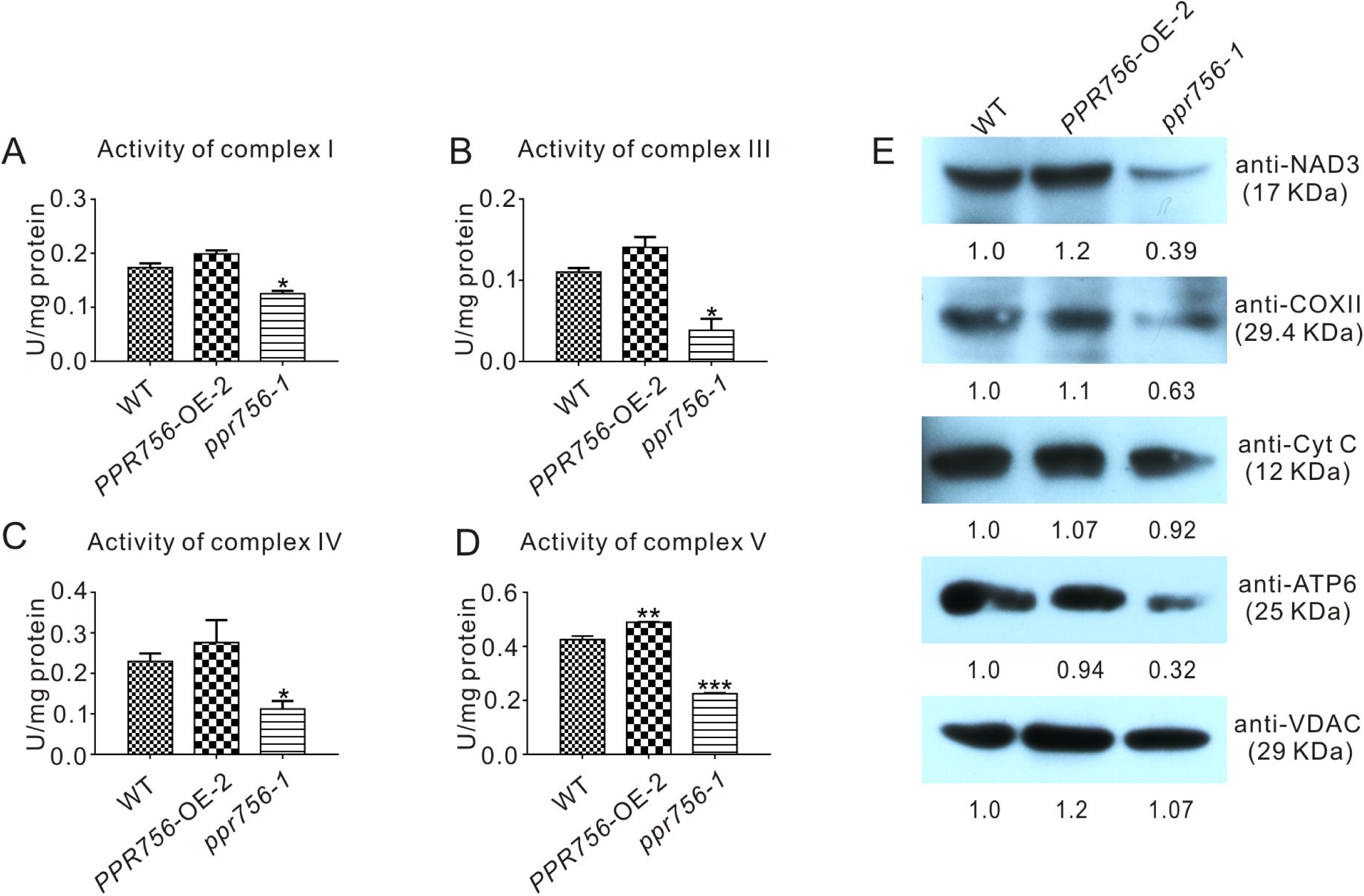
Detection of the activities and proteins in mitochondrial electron transport chain complexes. **(A–D)** In-solution determination of the activities of complex I, III, IV, and V in every 1 mg protein to response to their substrate in 1 μmol per minute. Bars represent the SDs from three independent biological replicates. Quantitative data were means ± SD based on three independent experiments (Student’s t-test; *, *P* < 0.05; **, *P* < 0.01; ***, *P* < 0.001). **(E)** Western blot detection of the components content in mitochondrial electron transport chain complexes. NAD3, COXII, and ATP6 represent complex I, complex IV, and complex V, respectively, and Cyt C with VDAC served as a control. The values of the samples were calculated by ImageJ.

**FIGURE 7.**
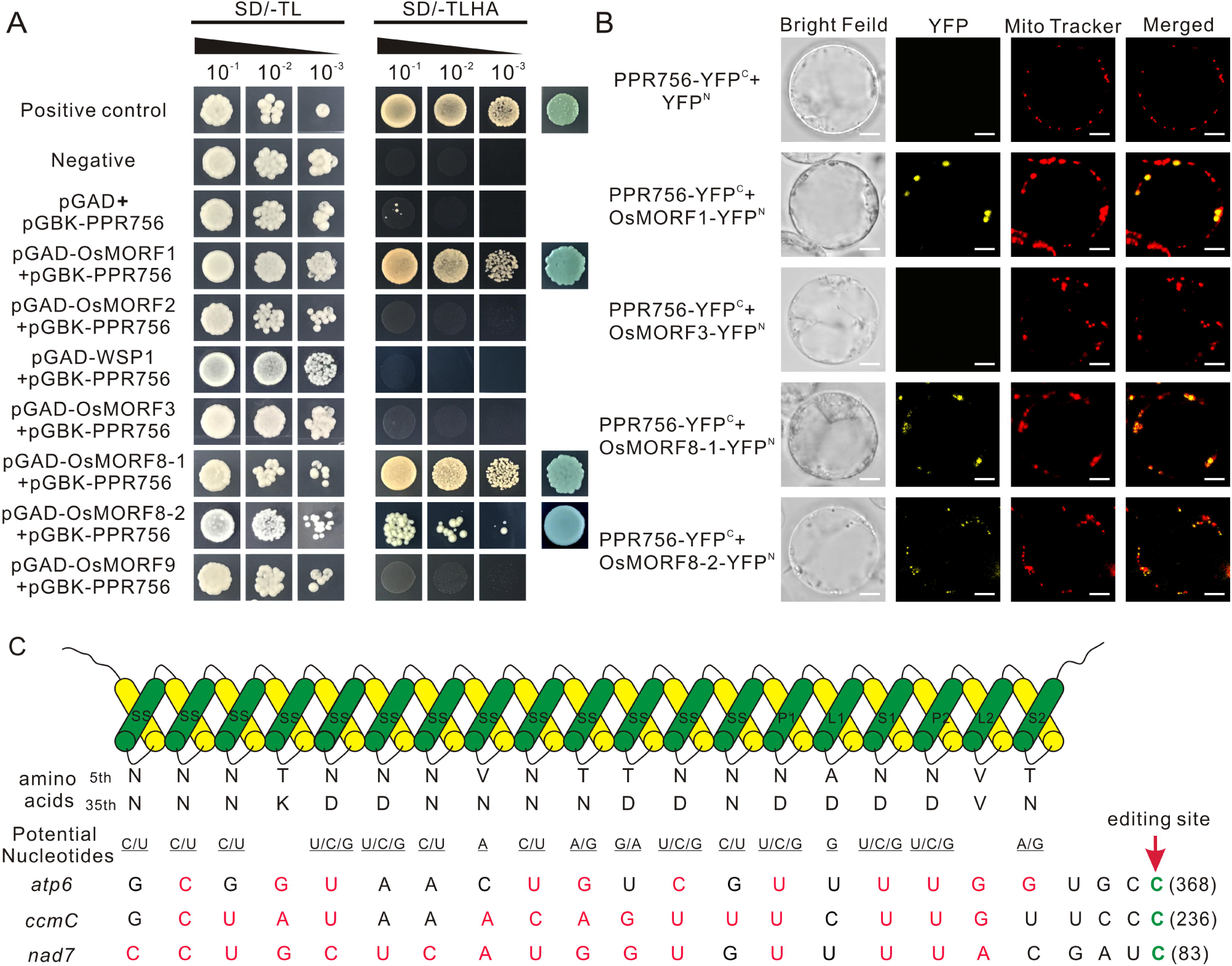
PPR756 directly interacts with OsMORF1 and OsMORF8s, and the PPR-RNA recognition model. **(A)** Yeast two-hybrid assays. PPR756 was cloned into the bait vector pGBKT7, and OsMORFs were cloned into the prey vector pGADT7. Transformants were grown on SD/-Trp/-Leu and SD/-Trp/-Leu/-His/-Ade dropout plates. The 53-T and 53-Lam interactions were used as positive and negative controls, respectively. **(B)** Bimolecular fluorescence complementation (BiFC) assays were performed to confirm the interactions between PPR756 and OsMORFs in rice protoplasts. The mitochondria were indicated by MitoTracker Red. Bars, 5 μm. **(C)** Potential recognition mechanism of PPR756 to its targets. The amino acid combination of position 5 and 35 in each PPR motif of PPR756 were aligned to the nucleotide upstream of the editing site. The potential recognized nucleotides are underlined and permissible matched nucleotides are colored in red, as well as the edited sites C are colored in green.

### Expression pattern of PPR756

To investigate the expression pattern of PPR756, bioinformatic data were first analyzed using the RiceXPro (http://ricexpro.dna.affrc.fo.jp), and results indicated that PPR756 was constitutively expressed in all the rice tissues during the whole development process but a higher level in the reproductive stage (Supplementary Figure 4). To confirm the expression profile, we carried out quantitative real-time PCR (qRT-PCR). PPR756 was expressed in all the tissues, including seedling, root, stem, leaf, young panicle, and mature spikelet. The relative expression of *PPR756* was much higher in mature panicles and seedlings, implying that it plays essential roles in these tissues (Figure 3C).

To characterize the expression pattern of PPR756 at the tissue level, we also generated the transgenic plants that harbor the p*PPR756*:GUS under the background of ZhongHua11. Different tissues of the transgenic plants were isolated for GUS histochemical staining. The staining results displayed that PPR756 was preferentially expressed in germinated seeds, seedlings, stems, and anthers (Figure 3D), which is consistent with the results of RT-PCR above. These findings demonstrate the important function of *PPR756* in seedlings, which affects the chlorophyll content of leaves, and its indispensable roles in the reproductive organs, which leads to sterile pollen.

### PPR756 affects multiple C-to-U RNA editing in mitochondria

The PLS-PPR proteins have been reported to typically function as site-specificity factors in RNA editing (Barkan and Small, 2014). According to the sub-localization results of PPR756, we checked all well-known RNA editing sites, 21 in the chloroplasts and 491 in the mitochondria, (Corneille et al., 2000;Notsu et al., 2002) via RT-PCR and direct sequencing. In the chloroplast, results showed no obvious RNA editing efficiency difference among the KO mutants, WT and the OE lines, which implies PPR756 does not function in the chloroplast directly. However, in the mitochondria, the RNA editing efficiency on three editing sites, *atp6-368, ccmC-236*, and *nad7-83*, were reduced in the RNAi line and KO mutants compared with the WT and OE lines, while no changes on other RNA editing sites were observed. (Figure 4 and Supplementary Figure 6 and 7). All of these three genes were involved in mitochondrial electron transport chain (ETC) complexes. The editing caused by the PPR756 protein led to amino acid changes: CCA(Pro)-to-CTA(Leu), CCA(Pro)-to-CTA(Leu), and TCA(Ser)-to-TTA(Leu) in *atp6, ccmC*, and *nad7*. Most editing failures generally influenced the function of the proteins as subunits of their corresponding complexes (Kim et al., 2009;Toda et al., 2012;Liu et al., 2013;Li et al., 2014;Xiao et al., 2018b), while a few were silent mutations (Zhu et al., 2012;Xiao et al., 2018a).

To investigate whether the transcript levels were affected by the editing deficiency, we used qRT-PCR to detect the relative mRNA levels of these targets of PPR756. The results demonstrated that there was almost no influence at the mRNA levels of the three targets (Figure 4B). Taken together, data suggest that PPR756 is responsible for the three RNA editing sites in mitochondria.

### PPR756 protein can directly bind to its targets both in vitro and in vivo

As previous studies reported, PPR protein could bind specifically to the surrounding RNA sequences of the editing sites as a trans-factor (Barkan et al., 2012;Nakamura et al., 2012). A series of experiments have also shown that the cis-element as a binding upholder was about 25 nucleotides of the upstream and 10 nucleotides of the downstream sequence surrounding the target editing sites (Okuda et al., 2006). To confirm PPR756 actually bound to its target transcripts, we generated recombinant PPR756 proteins. We initially failed to express the complete PPR756 with 18 motifs. Then, we removed the signal peptide and first S2 motif (residues 1–40) for expression, creating MBP-PPR756^41-756^-HIS protein, which contained the MBP tag in its N-terminus and 6xHis tag in the C-terminus (Supplementary Figure 8A). Recombinant tagged protein containing the two tags (MBP/HIS) only was used as a negative control. The two recombinant proteins were expressed in *Escherichia coli* and purified. The purified recombinant proteins were tested by Western blot using a His-tag antibody (Supplementary Figure 8B). The binding cis-element sequences of PPR756’s three targets were chosen by the −25 to +8 nucleotides surrounding the editing site according to previous research and labeled with fluorescent FAM. As a control, probe C (*nad5-1580*) was synthesized in our previous study, and it has been reported as the target of MPR25, another PPR protein in rice. To carry out the RNA electrophoresis mobility shift assays (REMSA), the purified recombinant proteins were dialyzed to remove the RNase contamination and incubated with the FAM-labeled RNA probes. The binding efficiency could be detected by the shift difference between the protein–RNA complexes and the free RNA probes. The retarded bands were observed when the targets were incubated with MBP-PPR756^41-756^ -HIS protein, while the tagged proteins could not bind to the RNA probes (Supplementary Figure 8C). No electrophoresis shift was observed when the probe C was incubated with MBP-PPR756^41-756^-HIS protein as a negative control, which suggested that PPR756 bound to its target RNAs specifically. We then performed the competitor assays using the corresponding unlabeled RNA probes to further confirm the preference of PPR756 for its targets. The binding capacity to the labeled probes gradually decreased following the increased competitor concentration, suggesting that PPR756 bound to these targets directly (Supplementary Figure 8D). To validate the binding activity in vivo, we next conducted RNA immunoprecipitation (RIP) in the OE transgenic plants to check the transcripts incorporated to PPR756. In this procedure, we used the Flag antibody to obtain the RNAs bound to PPR756, and the extracted RNA were converted to cDNA by reverse transcription (Figure 5B). Several negative controls were designed, and the Western blot was conducted to verify the RIP efficiency. Results showed the antibody anti-Flag could specifically capture PPR756-Flag, and the outputs were appropriate for further analysis (Figure 5A). Subsequently, we applied qRT-PCR using relevant primers to make sure of the proportion of these three target transcripts. The results showed that *atp6, ccmC*, and *nad7* transcripts displayed different binding efficiencies to PPR756; nevertheless, the negative control GAPDH did not (Figure 5C). The percentage ratios of these three targets were 6.014, 0.210, and 0.090, suggesting that PPR756 preferentially bound to *atp6* compared with other two targets (Figure 5D). This implies that PPR756 might bind to these via a different mechanism. Taken together, our results showed PPR756 could bind to the mitochondrial transcripts *atp6, ccmC*, and *nad7* directly and specifically both in vitro and in vivo.

### Dysfunction of PPR756 affects the activity of mitochondrial electron transport chain (ETC) complexes

Transcripts of *atp6* and *nad7* encode the subunit of respiratory complex V and I, respectively, and *ccmC* is involved in the biogenesis of cytochrome *c*, which transfers electrons from complex III to complex IV. According to the indispensable roles of *atp6, ccmC*, and *nad7* in theses complexes, we attempted to detect the activities of complexes of ETC in the WT, OE, and KO mutant lines. The in-solution determination results showed that in the KO mutant, the activity of complexes I, III, IV, and V were all reduced strikingly compared with the WT (Figure 6A–D). The data from OE transgenic lines showed that the activity of complexes I, III, IV, and V were increased, which was consistent with our expectations. Therefore, we speculated that these translated proteins without proper editing at the RNA level might reduce their biological functions to varying degrees. Several representative antibodies for ETC complexes were next employed. The results suggested that except for the slight reduction of cytochrome c (Cyt C), the others such as NAD3, COXII, and ATP6 all showed obvious decreases in the KO mutant compared with those in the WT (Figure 6E). This suggested that the decreased protein accumulation of these proteins in ETC complexes resulted in the reduction of the activities. The slight effect on Cyt C might have resulted from the defective editing site being far away from its functional WWD domain (Ahuja and Thony-Meyer, 2003). All these data indicated that dysfunction of PPR756 led to abortive editing and then induced the decrease of mitochondrial electron transport chain complexes’ activities.

### PPR756 interacts with OsMORF1 and OsMORF8s

Previous studies reported that some PPR proteins function with MORFs (multiple organellar RNA editing factors) as an editosome (Bentolila et al., 2012;Shi et al., 2016). Here, to make it clear whether PPR756 cooperated with MORFs, we investigated the interactions in vitro and in vivo. All of the seven MORFs in the rice database were cloned previously, including one MORF1-like (Os11g11020), two MORF2-like (Os04g51280, Os06g02600), one MORF3-like (Os03g38490), one MORF9-like (Os08g04450), and two MORF8-like (Os09g04670, Os09g33480). Yeast two-hybrid system was first used to determine the interactions. Interactions between PPR756 and OsMORF1/8s were detected, while no interaction between PPR756 and OsMORF2/3/9 was observed, indicating that PPR756 interacted with OsMORF1/8s but not with OsMORF2/3/9 (Figure 7A). Studies showed OsMORF8s were dual-localized in the mitochondria and chloroplasts, while OsMORF1 and OsMORF3 were sub-localized in the mitochondria only (Takenaka et al., 2012;Hartel et al., 2013;Glass et al., 2015;Xiao et al., 2018b). Therefore, the bimolecular fluorescence complementation (BiFC) was conducted to detect the interactions between PPR756 and all mitochondria-located OsMORFs in vivo. Yellow fluorescence signals were clearly observed when the PPR756 and MORF1 or MORF8s fusions were coexpressed in the rice protoplast, whereas no fluorescence was detected in PPR756 and OsMORF3, or the negative control, indicating the special and physical interactions between them in vivo (Figure 7B). Taken together, we demonstrated that PPR756 directly interacted with OsMORF1/8s in mitochondria, consistent with fact that the editing events of mitochondrial genes were affected rather than the chloroplast genes.

## DISCUSSION

### PPR756 is required for plant development

In plants, RNA editing is a familiar event that occurs in organelle transcripts, which is critical for gene repair and essential for the development of plants (Takenaka et al., 2013;Oldenkott et al., 2019). There are several types of RNA editing in plants including C-to-U, U-to-C, and A-to-I. The three editing events have different occurrences: C-to-U editing occurs in mRNAs and tRNAs, A-to-I editing in tRNAs, and U-to-C editing in mRNAs of a few non-flowering land plants (Chateigner-Boutin and Small, 2010). Previous study of the rice transcriptome revealed 21 and 491 C-to-U editing sites in the chloroplasts and mitochondria, respectively (Corneille et al., 2000;Notsu et al., 2002). Despite the mechanism and evolutionary significance of RNA editing still being obscure, several RNA editing factors have been characterized, such as PPRs, MORFs (also known as RIPs, RNA editing factor interacting protein), ORRM (organelle RNA recognition motif-containing), PPO1 (protoporphyrinogen IX oxidase 1), and OZ1 (organelle zinc finger 1) (Sun et al., 2016).

Up to now, approximately 65 nuclear encoded genes have been characterized for RNA editing in mitochondria and/or chloroplasts in Arabidopsis, rice, maize, and *Physcomitrella patens*, and so on. Only five of them, OGR1, OsSMK1, MPR25, OsPGL1, and OsPPS1, were identified in rice (Kim et al., 2009;Toda et al., 2012;Li et al., 2014;Xiao et al., 2018a;Xiao et al., 2018b). Here, we characterized a novel PPR gene, *PPR756*, which is involved in the RNA editing of *atp6*-*368, ccmC*-*236*, and *nad7*-*83*. NAD7 is the subunit of respiratory complex I, and ATP6 is the subunit of respiratory complex V. Therefore, the activities of complex I and V are expected to reduce, which may then influence the activities of complex III and complex IV. All of them are involved in energy production, and consequently, like most PPR mutants, loss of function of PPR756 led to defects of plant pleiotropic phenotypes. Furthermore, in this study, we showed that PPR756 expressed in most tissues but highly expressed in seedlings and anthers, exhibiting a spatial expression pattern. The pollen was sterile, and seed setting was extremely low compared to the WT, which was consistent with the high expression of PPR756 in anthers. We speculated that the energy for seedlings and anthers is in great demand, any shortage of which could lead to developmental defects in the corresponding stage.

### PPR756 shares the PPR-RNA recognition model

A commonly held view is that PPR proteins can directly bind to target RNA in a specific manner, which is confirmed by bioinformatic and structural analysis (Barkan et al., 2012;Yin et al., 2013). A recognition model has been verified that one PPR motif corresponded to one nucleotide (Barkan et al., 2012;Okuda et al., 2014;Kindgren et al., 2015), in which a di-residues combination was especially important. The di-residues were the 5th and 35th residue in one motif (Yin et al., 2013), which were also named residues 6 and 1’ in Barkan *et.al* and residues 4 and ii in Yagi *et.al* (Barkan et al., 2012;Yagi et al., 2013)

Most PLS-PPR proteins consist of P, L, and S motifs in turn, and usually with the pattern (P1-L1-S1)_n_-P2-L2-S2 in front of the C-terminus (Rivals et al., 2006).. However, in this study, PPR756 was distinctive, including numbers of continuous SS motifs in front of P1-L1-S1 -P2-L2-S2. To evaluate the PPR–RNA recognition model, we analyzed the associations between PPR756 and its three target genes on the basis of the reported general recognition mechanism (Yan et al., 2019). We found that the match indices between PPR756 and its targets *atp6, ccmC*, and *nad7* were 11/19, 14/19, and 16/19, respectively (Figure 7C), with such relatively high match scores suggesting that PPR756 could bind its target genes directly in accordance with the PPR–RNA recognition model. A more interesting finding in the match alignments showed a highly matching score from the last 8 nucleotides (−4 to −12), implying that the latter might be more important than the former during recognition. In addition, the secondary structures of target RNA may also have effects on the binding affinities of PPR proteins (Williams-Carrier et al., 2008;Prikryl et al., 2011;Kindgren et al., 2015;Miranda et al., 2017;McDermott et al., 2018).

One of the most essential issues is the deaminase activities during RNA editing events. Given that PPR756 belongs to PLS-E subgroup without DYW domain, we speculate that other unknown factor with deaminase activity is needed. Previous studies showed that some DYW proteins acted as indispensable partners in hundreds of editing sites in Arabidopsis, which were consisted of few PPR motifs and DYW domain without canonical E and E+ domains (Boussardon et al., 2012;Andres-Colas et al., 2017). They could be recruited by other PLS-subclass PPR proteins without DYW domain. For example, DYW1 was essential for the editing of *ndhD-1* site via interacting with another PLS-E protein, CRR4 (Boussardon et al., 2012). Recent studies reported that DYW2 can interact with several PLS-E+ PPR proteins to regulate many RNA editing sites in Arabidopsis (Andres-Colas et al., 2017;Guillaumot et al., 2017;Malbert et al., 2020). However, none of PPR proteins without DYW motif was characterized in rice so far, therefore, we speculated that PPR756 could function in an editosome associated with other PPR proteins, and MORFs.

## CONCLUSION

Our results suggested that PPR756 can recognize and bind three mitochondrial gene *atp6, nad7* and *ccmC*, which was subunit of ETC (electron transport chain) complex V and I, and critical for the biogenesis of cytochrome *c* in rice, respectively as well as participated in the their RNA editing processes. Loss function of PPR756 caused the extremely lower editing efficiency of the three mitochondrial genes and further led the activity reduction of the ETC complexes. While PPR756 was abundant in the anther which may need more energy in the reproductive stage, the dysfunction of PPR756 then induced sterile pollens and abortive seeds. These results shed light on us a novel pentatricopeptide repeat protein and further enriched our comprehension of editing factors in rice.

## DATA AVAILABILITY

Sequence data from this work can be found in the GenBank databases under the following accession numbers: PPR756 (Os12g19260), MORF1 (Os11g11020), WSP1(Os04g51280), MORF2 (Os06g02600), MORF3 (Os03g38490), MORF8-1 (Os09g33480), MORF8-2(Os09g04670), MORF9 (Os08g04450).

## AUTHOR CONTRIBUTIONS

Q.Z, J.Hu and Y.Z designed the experiments. Q.Z carried out most experiments. Y.X performed the subcellular location and BiFC experiments. J. Huang contributed to the protein purification and field management. K.Z performed the mitochondria isolation and RNA Immunoprecipitation experiments. H.X and X.Q performed the data analysis. L.Z contributed to the RNAi transgenic lines. Q.Z and J.Hu wrote the paper with feedback from all authors.

## ACKNOWLEDGEMENTS

We appreciate professor Yaoguang Liu (South China Agricultural University) for the gifts of pYLCRISPR/Cas9 vectors. We also appreciate professor Yunkuang Liang (Wuhan University) for constructive suggestions. We appreciate the contributions made by professor Yingguo Zhu (Wuhan University), and we would like to pay tribute to him in this article. This manuscript has been released as a pre-print at https://www.biorxiv.org/content/10.1101/844241v1(Zhang et al., 2019).

