## Supplemental figures for "The rice pentatricopeptide repeat protein PPR756 is involved in pollen development by affecting multiple RNA editing in mitochondria": Supplementary Data-0506.pdf

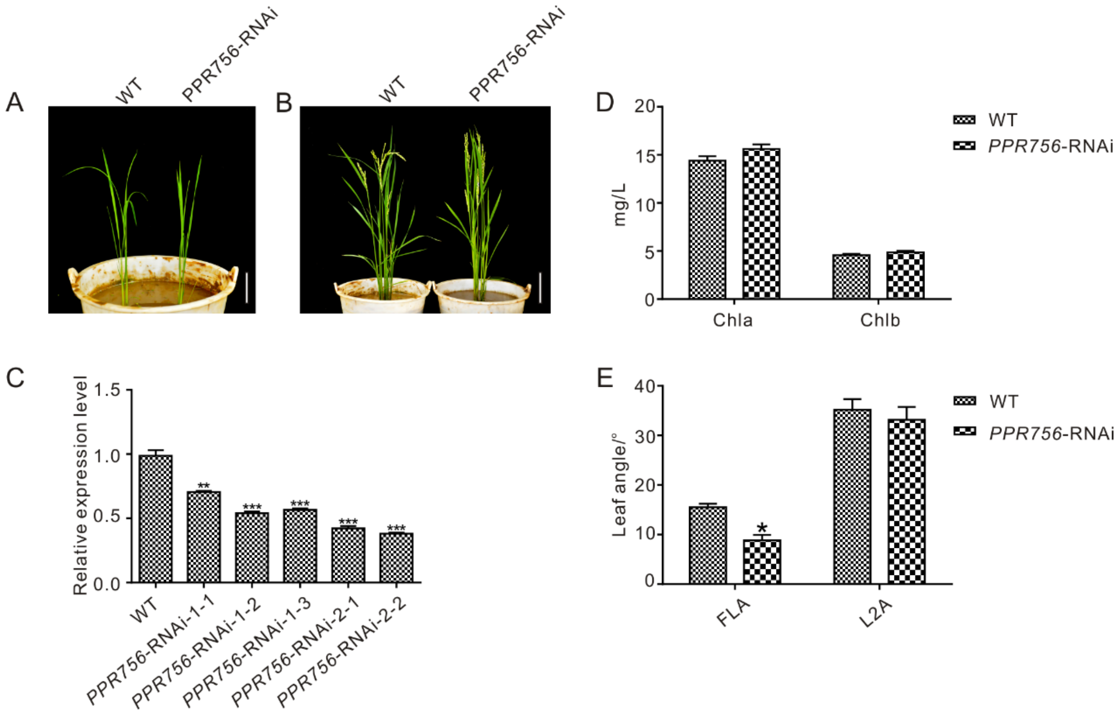

**Supplementary Figure 1.** Phenotypic characterization of *PPR756* RNAi lines.

**(A, B)** Plant phenotype comparison of the WT and *PPR756* RNAi lines at rice seedling and reproductive stage. Bars 5 cm for **(A)** and 10 cm for **(B)** respectively. **(C)** The relative expression level of *PPR756* transcript in *PPR756* RNAi lines were detected by real-time qPCR. **(D)** The chlorophyll a (Chla) and chlorophyll b (Chlb) content of leaves in the WT and series of independent *PPR756* RNAi lines. The unit of content is mg/L. **(E)** Comparison of leaf angles in the WT and *PPR756* RNAi lines. FLA, flag leaf angle, L2A, the last second leaf angle. The unit is "o". All quantitative data were means ± SD based on three independent experiments (Student's t-test; *, $P < 0.05$; **, $P < 0.01$; ***, $P < 0.001$).

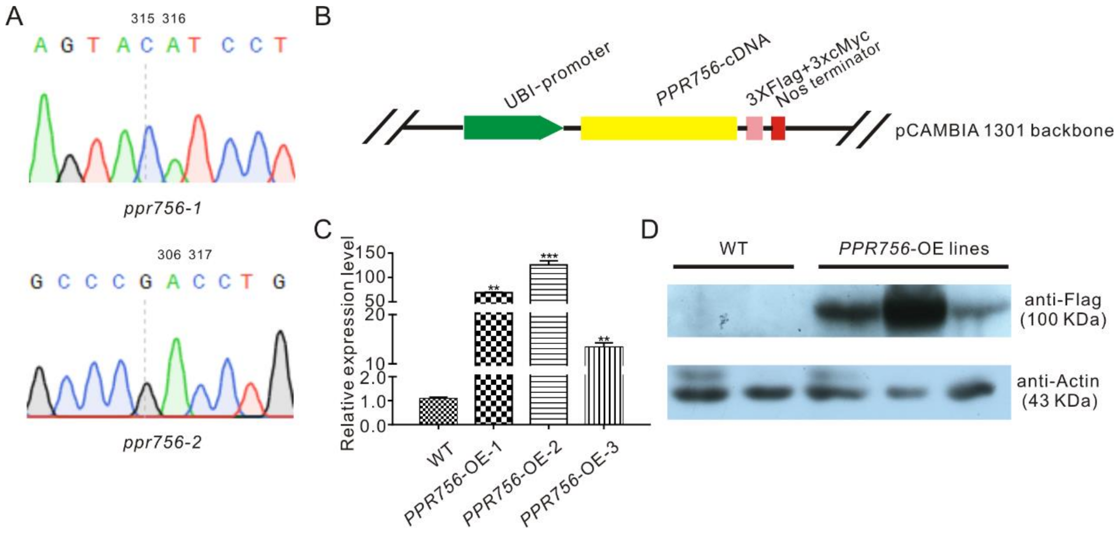

**Supplementary Figure 2.** Construction and validation of *PPR756* transgenic lines. **(A)** The sequencing analysis of *ppr756* mutants. **(B)** The structure scheme of *PPR756* overexpression construct. **(C)** The transcript level detection in the WT, three independent *PPR756* OE lines. Rice *actin* gene served as a control. **(D)** Immunoblot assays in the WT three independent *PPR756* OE lines. Actin was served as a control, and Flag signal represents the fusion protein PPR756-Flag-cMycs. Quantitative data were means ± SD based on three independent experiments (Student's t-test; *, $P < 0.05$; **, $P < 0.01$; ***, $P < 0.001$).

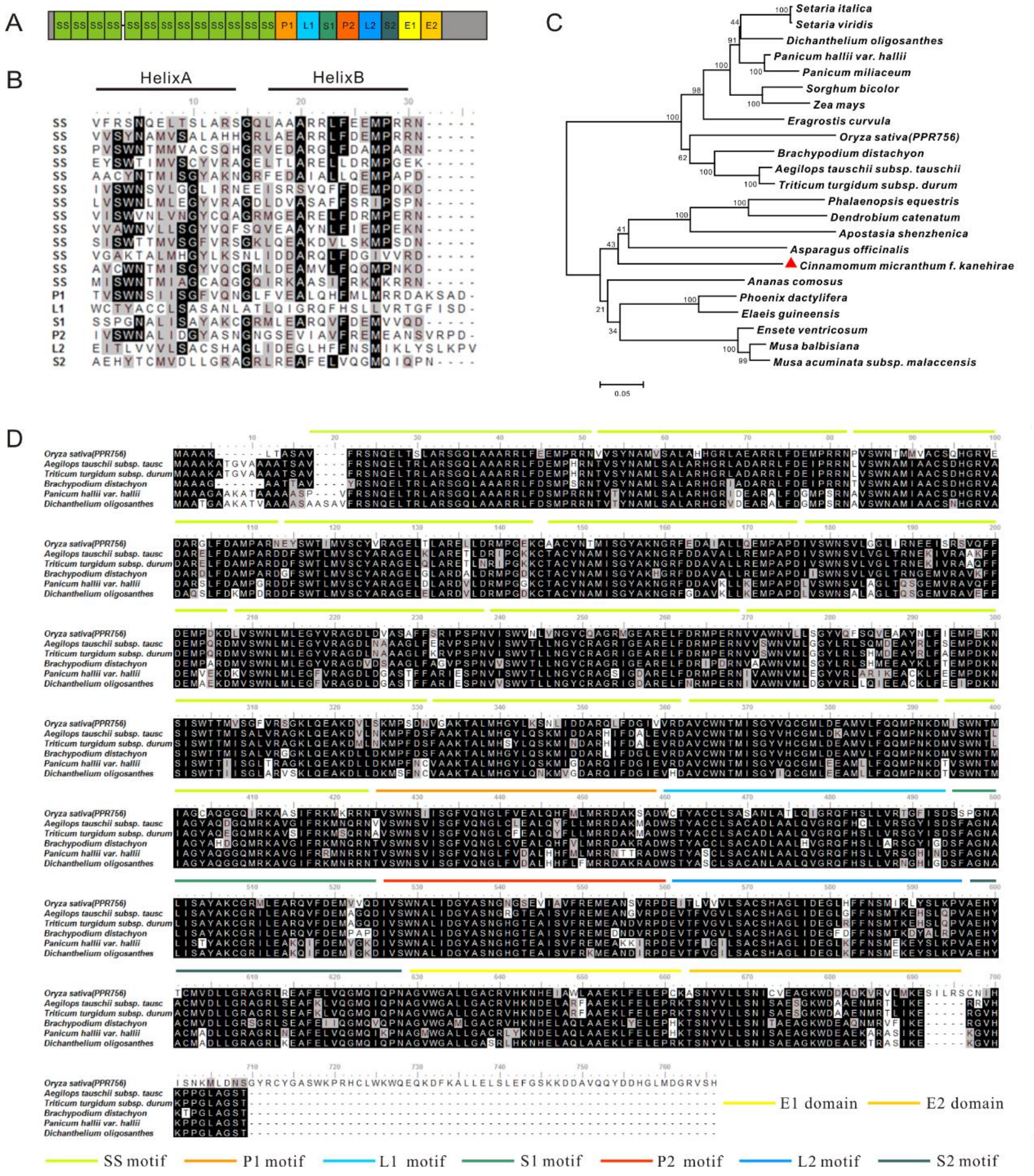

**Supplementary Figure 3.** Structure and sequence alignment of PPR756 protein.

**(A)** PPR and E motifs of PPR756 protein are indicated and marked respectively. **(B)** Amino acids sequence alignment of 19 PPR motifs of PPR756, highly conserved amino acids are marked in dark. **(C)** Phylogenetic tree analysis of PPR756 homologs. The analysis was constructed by MEGA 5.2 program and showed the homologs over 55% similarity including including *Aegilops tauschii*, *Triticum turgidum*, *Brachypodium distachyon*, *Panicum hallii*, *Dichanthelium oligosanthes*, *Setaria italic*, *Eragrostis curvula*, *Setaria viridis*, *Sorghum bicolor*, *Zea mays*, *Panicum miliaceum*, *Phoenix dactylifera*, *Ananas comosus*, *Ensete ventricosum*, *Cinnamomum micranthum f. kanehirae*, *Musa balbisiana*, *Asparagus officinalis*, *Musa acuminata subsp.*

*Malaccensis, Phalaenopsis equestris, Apostasia shenzhenica, Dendrobium catenatum* were almost monocots, and only one was a dicot (marked by triangle). **(D)** Detailed sequence alignment of rice PPR756 protein with some orthologs. The alignment showed very highly conservation among PPR756 and its orthologs. The different motifs are marked using various colors of straight lines.

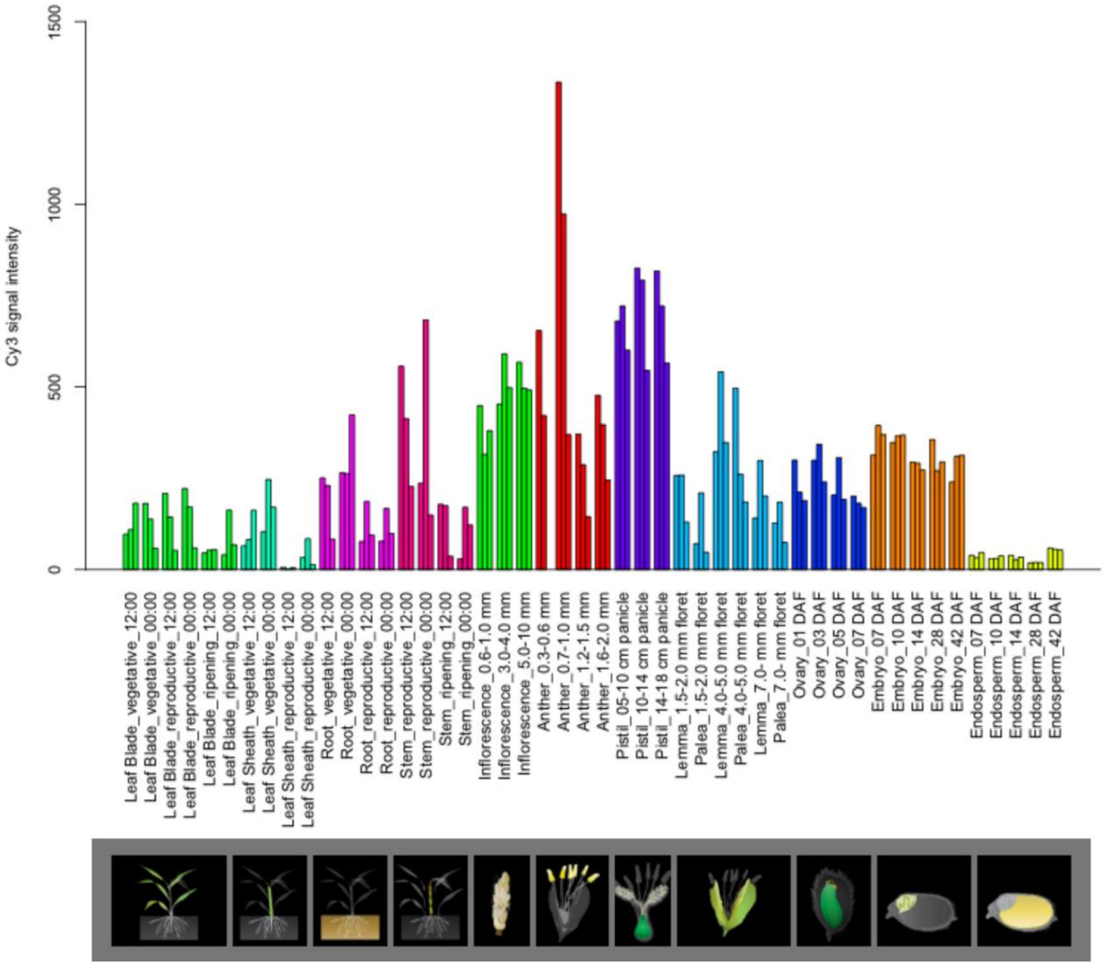

**Supplementary Figure 4.** Predicted spatio-temporal expression pattern of PPR756.

Predicted spatio-temporal expression pattern of PPR756 from the RiceXPro database.

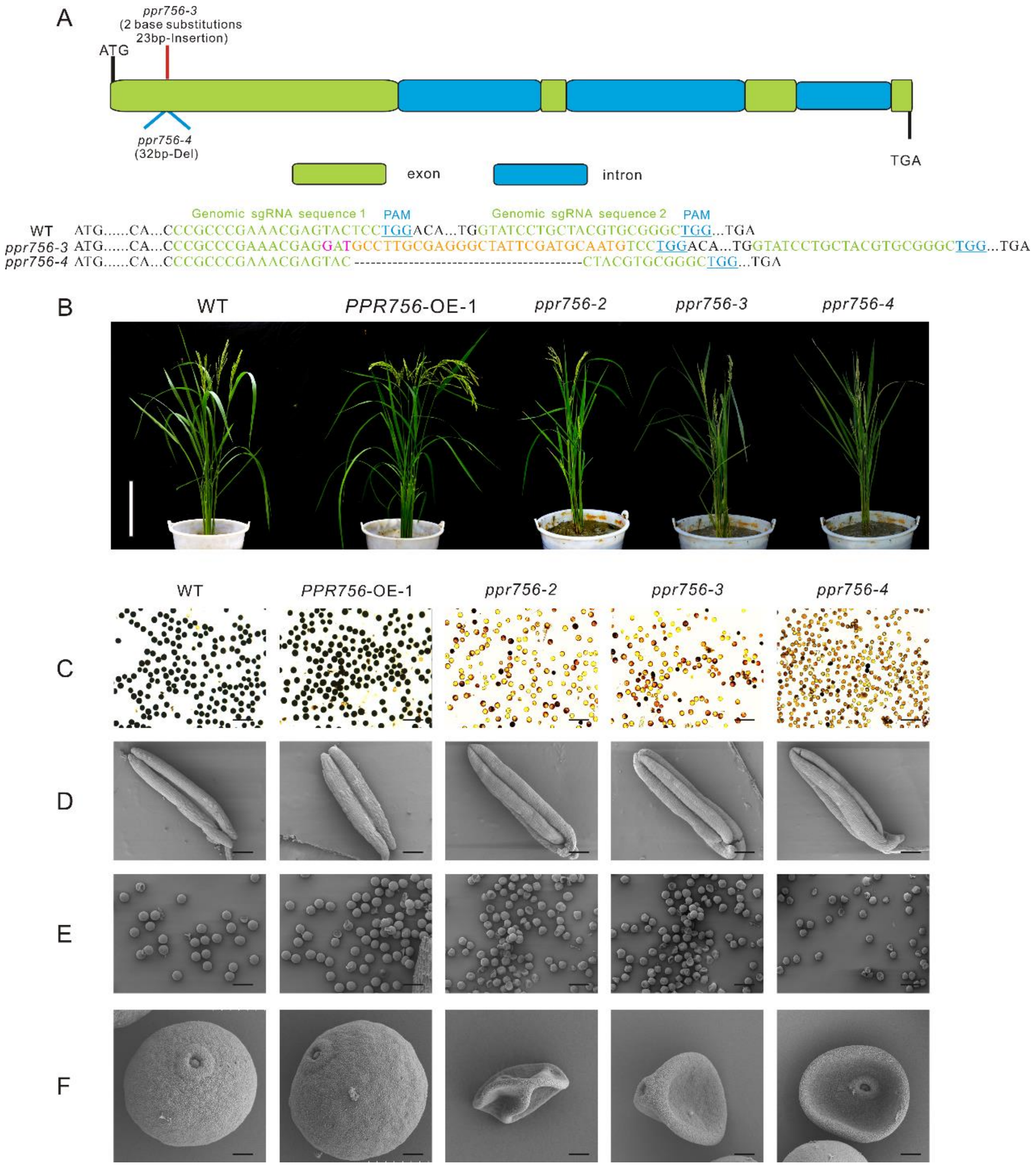

**Supplementary Figure 5.** Phenotypic characterization, fertility detection and scanning electron microscopy observations in the WT, *PPR756* OE line and *ppr756* mutant lines. **(A)** The structures of *PPR756* and *ppr756* mutants. Other *ppr756* mutant lines showed with 2 base substitutions 23 bp insertion in *ppr756-3* line and 32 bp deletion in *ppr756-4* line. **(B)** Plant phenotypes comparison of the WT, PPR756 OE line and *ppr756-2*, *ppr756-3*, *ppr756-4* mutant lines at reproductive stage. Bars, 20 cm. **(C)** Comparison of pollen fertility of WT, the *PPR756* OE line, and *ppr756* mutant lines by potassium iodide (1% $I_2$-KI) staining. The darkly stained pollen is fertile, whereas the lightly or non-stained pollen is sterile. Bars, 200 μm. **(D)** Scanning electron microscopy (SEM)

observation of anthers of WT, the *PPR756* OE line, and *ppr756* mutant lines. Bars, 200 μm. Representative results from three independent experiments. **(E, F)** Scanning electron microscopy (SEM) observation of pollens of the *PPR756* OE line and *ppr756* mutant lines. Bars, 100 μm for **(E)** and 10 μm for **(F)**.

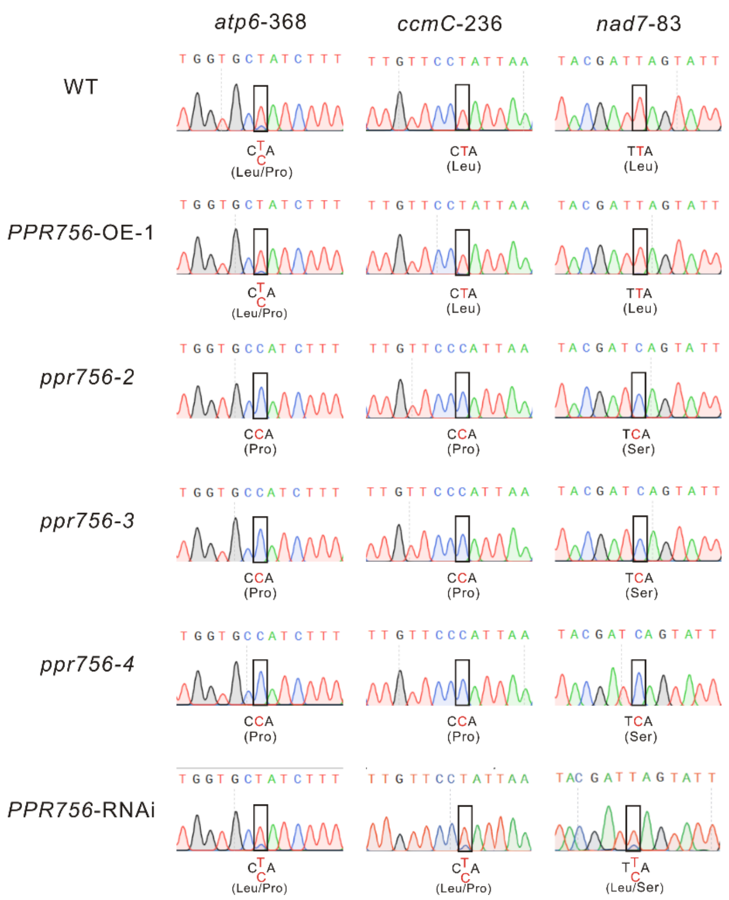

**Supplementary Figure 6.** RNA editing analysis in the WT, *PPR756* OE line, *ppr756* mutant lines and *PPR756* RNAi line.

RNA editing analysis of *atp6-368, ccmC-236, nad7-83* in the WT, *PPR756* OE line, *ppr756-2*, *ppr756-3*, *ppr756-4* mutant lines and *PPR756* RNAi line.

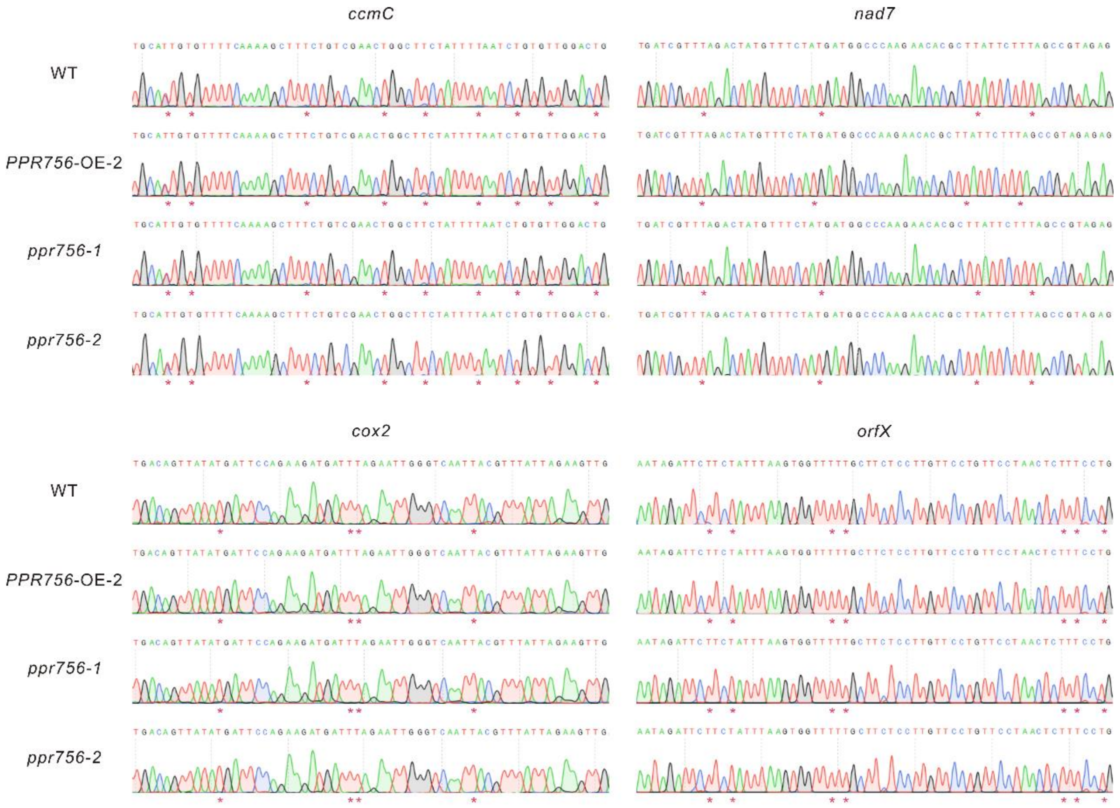

**Supplementary Figure 7.** RNA editing analysis of unaffected sites in the WT, *PPR756* OE line and *ppr756* mutant lines.

RNA editing analysis of unaffected mitochondrial edited sites in the WT, *PPR756* OE line and *ppr756* mutant lines. The edited sites are marked by “*”.

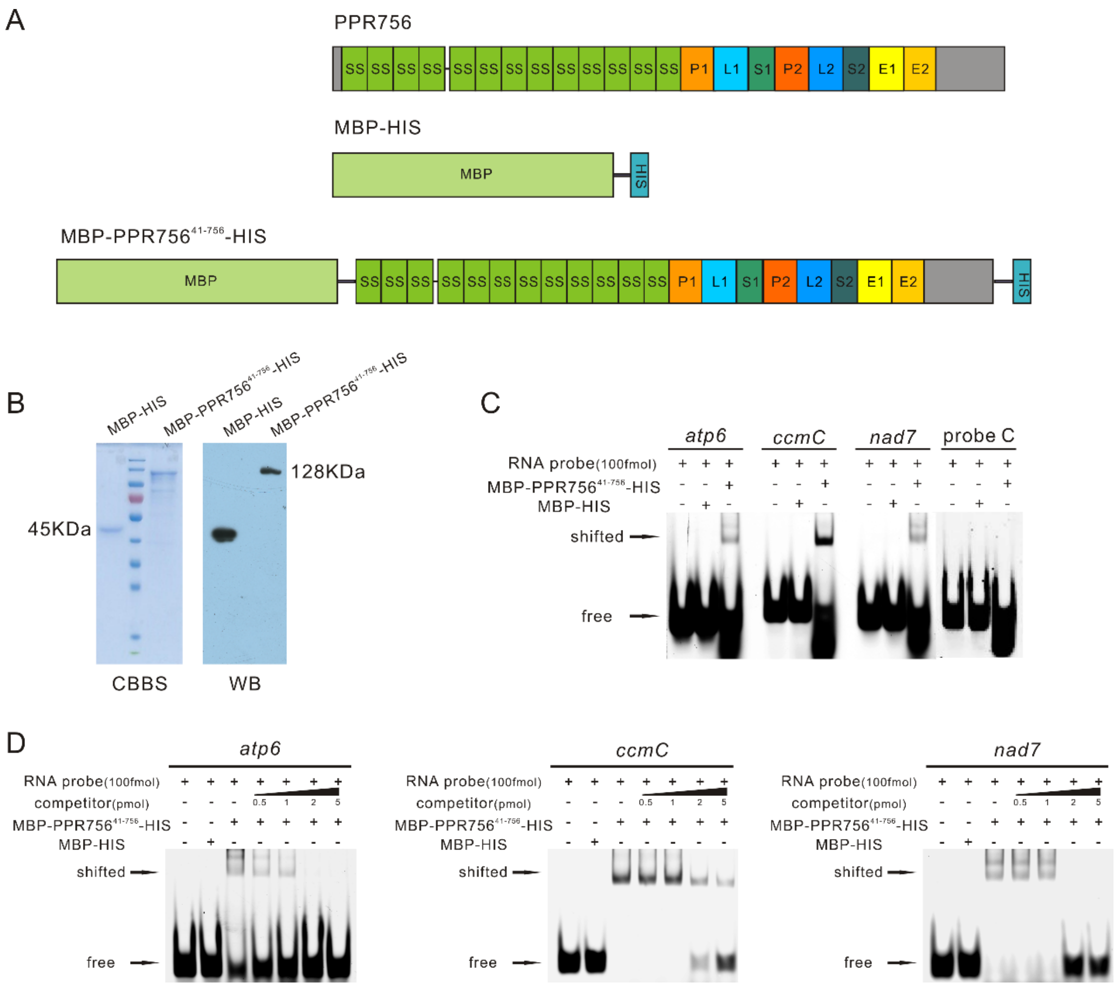

**Supplementary Figure 8.** PPR756 can directly bind to its target genes in vitro.

**(A)** Schematic structures of expressed recombinant proteins. **(B)** The expressed and purified recombinant proteins were validated by Coomassie brilliant blue staining (left) and Western blots (right). CBBS, Coomassie brilliant blue, WB, Western blot. **(C)** RNA electrophoresis mobility shift assays (REMSAs) of the recombinant MBP-PPR756$^{41-756}$-HIS protein and MBP-HIS tags with the FAM-labeled RNA probes. MBP-HIS tags and probe C served as a negative control. **(D)** The corresponding non-labeled RNA probes in a concentration gradient were used as competitors for the REMSA of MBP-PPR756$^{41-756}$-HIS protein with its target probes. MBP-HIS tags served as a negative control.

### Table S1 Primers used in this study

| Name | Sequences(5'-3') | Purpose |
|---|---|---|
| *PPR756*-RNAi-F | GGGGACAAGTTTGTACAAAAAAGCAGGCTGGAAGCTCACCGCCTCTGCTGTGTTTA | RNAi |
| *PPR756*-RNAi -R | GGGGACCACTTTGTACAAGAAAGCTGGGTC ACCAGCCCGCACGTAGCAGGAT | |
| *PPR756*-cDNA-F | ATGGCAGCTGCGAAGCTCAC | Amplification |
| *PPR756*-cDNA-R | ATGACTTACTCTGCCATCCATGAGC | |
| *PPR756*-RT-F | AGCGTAATGTTGTGGCTTGGAATGT | RT-PCR |
| *PPR756*-RT-R | GGAATACTGCTATCACCTCAGACCC | |
| *PPR756*-HBT-F | CTTGCTCCGTGGATCCGCCACCATGGCAGCTGCGAAGCTCAC | Subcellular localization |
| *PPR756*-HBT-R | TGCTCACCATGGATCCATGACTTACTCTGCCATCCATGAGC | |
| *PPR756*-qPCR-F | GCCTTGATTTCTGCCTAT | qRT-PCR |
| *PPR756*-qPCR-R | ACCTCAGACCCATTACCA | |
| *PPR756*$^{41\text{-}756}$-Pmal-2X-F | ATCCTCTAGAGTCGACAACGTGGTCTCCTACAACGCGAT | Prokaryotic expression |
| *PPR756*$^{41\text{-}756}$-Pmal-2X-R | TTGCCTGCAGGTCGACGCAGCCGATCTCAGTGGT | |
| *PPR756*-*BIFC-F* | CGCCACTAGTGGATCCGCCACCATGGCAGCTGCGAAGCTCAC | BiFC |
| *PPR756*-BIFC-R | TACTATCGATGGATCCATGACTTACTCTGCCATCCATGAGC | |
| *PPR756*-pro-F | GGAATTCCTCTATCCAAAACTGACAAACCAAGG | GUS expression |
| *PPR756*-pro-R | GGAATTCCTGCCGCCAGATACCACAA | |
| *PPR756*-OE-F | CGACTCTAGAGGATCGCCACCATGGCAGCTGCGAAGCTCAC | Overexpression |
| *PPR756*-OE-R | GTTTCTGCTCGGATCGCCACCATGACTTACTCTGCCATCCATGAGC | |
| *PPR756*-Y2H-F | GGCCATGGAGGCCGCCACCATGGCAGCTGCGAAGCTCAC | Yeast two-hybrid assays |

| | | |
|---|---|---|
| *PPR756*-Y2H-R | <u>GGCCTCCATGGCCAT</u>ATGACTTACTCTGCCATCCATGAGC | |
| *actin*-Q-F | CTTCATAGGAATGGAAGCTGCGGGTA | qRT-PCR |
| *actin*-Q-R | CGACCACCTTGATCTTCATGCTGCTA | |
| *atp6*-qpcr-F | ATTTGGTCTTGATATGGGTA | |
| *atp6*-qpcr-R | ATTTGGCACTGACTTTCC | |
| *ccmC*-qpcr-F | GAACAAATAGGTGGAAATG | |
| *ccmC*-qpcr-R | GAGCCAAAGTAATGAGAA | |
| *nad7*-qpcr-F | ACGGAGAAGTGGTGGAAC | |
| *nad7*-qpcr-R | GCTGAAGAATGAGCGTGT | |
| *nad5*-qpcr-F | GACAAGGGTGCTATTGAG | |
| *nad5*-qpcr-R | GAGTCCCACATACGAGAA | |
| *GAPDH*-QPCR-F | AAGCCAGCATCCTATGATCAGATT | |
| *GAPDH*-QPCR-R | CGTAACCCAGAATACCCTTGAGTTT | |
| *PPR756*-crispr-U6a-F | GCCGGTATCCTGCTACGTGCGGGC | CRISPR sytem |
| *PPR756*-crispr-U6a-R | AAACGCCCGCACGTAGCAGGATAC | |
| *PPR756*-crispr-U3-F | GGCACGCCCGAAACGAGTACTCC | |
| *PPR756*-crispr-U3-R | AAACGGAGTACTCGTTTCGGGCG | |
| *PPR756*-gRT-U6a+ | GTATCCTGCTACGTGCGGGCGTTTTAGAGCTAGAAAT | |
| *PPR756*-gRT-U3+ | CGCCCGAAACGAGTACTCCGTTTTAGAGCTAGAAAT | |
| *PPR756*-OsU6aT- | GCCCGCACGTAGCAGGATACGGCAGCCAAGCCAGCA | |
| *PPR756*-OsU3T- | GGAGTACTCGTTTCGGGCGTGCCACGGATCATCTGC | |
| CAS9-PCR-F | CCACTCCATCAAGAAGAACCTCATC | |
| CAS9-PCR-R | CGATAAGGAAGTGACCACGGAAC | |
| *PPR756*-TEST-F | GGTCTCCTACAACGCGATGGTCT | |
| *PPR756*-TEST-R | CACAACATTACGCTCAGGCATTCTA | |
| *atp6*-FAM | UAUGGCGGUAACUGUCGUUUUGGUGCCAUCUUUAU | RNA-EMSA |

| | | |
|---|---|---|
| *ccmC*-FAM | ACAGCUAUAAACAGUUUCUUGUUCCCAUUAACAA | |
| *nad7*-FAM | UCCUGCUGCUCAUGGUGUUUUACGAUCAGUAUUCC | |
| *nad5*-FAM | AAAACUAAUACCUAUUCUGUUUAGUACUUCAGGUG | |
| *atp1*-1-F | ATGGAATTCTCACCCAGAGC | Mitochondrial RNA editing assay |
| *atp1*-1-R | TATAGGAACCAGGCTATCCA | |
| *atp1*-2-F | TAGAAAGAGCCGCTAAAC | |
| *atp1*-2-R | CTAATTAATCTCCTTCGCAG | |
| *atp6*-1-F | AATTACTCATTTTAATGGAG | |
| *atp6*-1-R | CCACTGCCATTAGCACCTTT | |
| *atp6*-2-F | GAAAATGACTTGTCACTGTG | |
| *atp6*-2-R | TTTCGATTACAATCATGTGG | |
| *atp9*-F | GCAAAGTCAAGTCTCCACGA | |
| *atp9*-R | CAAAGAGAGATATCTACACC | |
| *ccmB*-F | ATGAGACGACTCTTTCTTGA | |
| *ccmB*-R | TCAATCTTGTAAACTAATCG | |
| *ccmC*-F | ATGTCAGTTTCGTTATTACA | |
| *ccmC*-R | CTAGGTTTTTAGTGGTATTC | |
| *ccmFc*-1-F | GGTCCAACTACAGAACTTCT | |
| *ccmFc*-1-R | AGTAGTCGTGACCAACAGCCA | |
| *ccmFc*-2-F | TGTTGGTCACGACTACTACAAAAAA | |
| *ccmFc*-2-R | TCAATTCCAATGCAACTTAT | |
| *ccmFn*-1-F | ATGTCTATAAATGAATTTTC | |
| *ccmFn*-1-R | AACCACGGGAGCGCCCAGCG | |
| *ccmFn*-2-F | TTGACGGAGCTCTTGCCATT | |
| *ccmFn*-2-R | CGAGCTTCTTATATGGGATC | |
| *ccmFn*-3-F | CAGGACCAGGAACCAATTCG | |
| *ccmFn*-3-R | CTACGGACGGGACGAAATCC | |
| *cob*-1-F | ATGACTATAAGGAACCAACG | |
| *cob*-1-R | AGCCAGATGAAGAAGACTGG | |
| *cob*-2-F | CATTGGGTGTACATTCTGAG | |
| *cob*-2-R | CTAAGAGACTGATCCGGTGC | |
| *cox*1-F | AATGCTCTGAGCAGTTTCGG | |
| *cox*1-R | CTAGCTTTTTGTCTCTTTGA | |
| *cox2*-F | GTATAGTAGTCTCATTGGCC | |
| *cox2*-R | TCTTTCAAAGTCACCGCTTC | |
| *cox3*-F | TATTACCAAGCACCCTCCAC | |
| *cox3*-R | TCATATACCTCCCCACCAAT | |
| *nad1*-1-F | AGGCCCGATCATGAGTGAAT | |
| *nad1*-1-R | GAAAATGATCTGGTTGGACG | |
| *nad1*-2-F | CGTCCAACCAGATCATTTTC | |

| | |
|---|---|
| *nad1*-2-R | TCATATTGGCATACTCTCCC |
| *nad1*-3-F | GGGAGAGTATGCCAATATGA |
| *nad1*-3-R | TTAAGGGAGCCATCGAAAGG |
| *nad2*-1-F | CCCACTTCGATCAATTAGCC |
| *nad2*-1-R | CGCTATATATTTGACACGGG |
| *nad2*-2-F | GCTCTAGCCAAAACGAATCC |
| *nad2*-2-R | CCGAACAATAATATTCCAGAG |
| *nad3*-F | ATGGACAACATTTTTTTTGG |
| *nad3*-R | TTACTCCCAATCCAAAGCAC |
| *nad4*-1-F | TAGAACATTTCTGTGAATGC |
| *nad4*-1-R | CCCATACCCCTATAATGATG |
| *nad4*-2-F | GCCCATATGAATTTGGTGAC |
| *nad4*-2-R | CACTAAGTTACTTACGGATG |
| *nad4L*-F | ATGGATCTTATAAAATATTT |
| *nad4L*-R | TTAACCTTGAATGCAATTTA |
| *nad5*-F | GGGAGTCTCTTTGTAGGATA |
| *nad5*-R | TAAGAAAACTGCTCACTAAC |
| *nad6*-F | ATGATACTTTCAGTTTTGTC |
| *nad6*-R | TTAGTAAATCGTGATTTGGT |
| *nad7*-1-F | TGACGACTAGGAACGGGCAA |
| *nad7*-1-R | CCACTAATCGTTGTTTCCAG |
| *nad7*-2-F | CACAGCAAGCAAAGGATTGG |
| *nad7*-2-R | CTATCTATCTACCTCTCCAAACACA |
| *nad9*-F | ATGGATAACCAATCCATTTT |
| *nad9*-R | TTATCCGTCGCTACGCTGTT |
| *orf25*-F | ATGGGATTGAGTTCAACGGA |
| *orf25*-R | CTTTCACTCACTGAAAAGTG |
| *OrfB*-F | ATGCCTCAACTTGATAAATT |
| *OrfB*-R | TTAGATTATGCTTCCTTGCC |
| *OrfX*-F | GCCGAAAATGCATTTATCCT |
| *OrfX*-R | CCACAAAGATAGCAAACTCG |
| *rpl16*-F | ATGGAAAAACATCTTGTAAT |
| *rpl16*-R | TTACGACCACTGAACAAACT |
| *rps11*-F | ATGCCTCAGGAAAAAAACAACG |
| *rps11*-R | TCACGATCCCGGTAGAGGGA |
| *rpl2*-F | ATCCAGGTCAAGGCGCAAAG |
| *rpl2*-R | TTTCTAAGCTTACGTGCACC |
| *rpl5*-F | ATGTTTCCACTCCATTTTCA |
| *rpl5*-R | GATCGAAACGACTTTCCTGC |
| *rps1*-F | ATGTTCTTGGTGGATGCAGG |
| *rps1*-R | TCAAGTTCTTGTTTGATCTG |
| *rps2*-F | TGAAAAAGACCAATCAAATCAAACT |

| | | |
|---|---|---|
| *rps2*-R | GGGTTCGTGCACAGATTTAC | |
| *rps3*-1-F | GGCACGAAAAGGAAATCCAA | |
| *rps3*-1-R | TATTAAAAAAGTATTGCATG | |
| *rps3*-2-F | CTCTTTCCTTTCTTCGGTGC | |
| *rps3*-2-R | ATTTCGTACGTTTCGGATATAGCAC | |
| *rps4*-1-F | ATGCCTGCATTAAGATTTAA | |
| *rps4*-1-R | AGTTGTGAGTAAGCGGAACC | |
| *rps4*-2-F | GCGGAAAACCGAAAAAGAGC | |
| *rps4*-2-R | TTATATGTTTTGGCCACGTC | |
| *rps7*-F | ATGGGGGACTTTGATGGTGA | |
| *rps7*-R | TTACCACCATCTGAAATGCG | |
| *rps13*-F | ATGTTATATATCTCAGGAGC | |
| *rps13*-R | TCATTTCCGAATTAGCTTGC | |
| *rps19*-F | ATGCCACGACGATCTATATG | |
| *rps19*-R | TTACTTTTTCCCCTTTCTGC | |
| *atpA*-F | CCCAGGGGATGTTTTTTATT | Chloroplast RNA editing assay |
| *atpA*-R | TGAAAAAAGCGTCCATTGTC | |
| *ndhA*-F | ATGATAATAGACAGGGTACAGG | |
| *ndhA*-R | TTATAGTGAAACAAGTTGGGAAG | |
| *ndhB*-F | ATGATCTGGCATGTAC AGAATG | |
| *ndhB*-R | CTAAAAGAGGGTATCCTGAGCA | |
| *ndhD*-F | ATTTTGGCTTCCTTATTGC | |
| *ndhD*-R | GCCTCTACCCTGTCAACG | |
| *ndhF*-F | ATATGCATGGGTAATCCCTC | |
| *ndhF*-R | AGTGGCTCCTAAGAAAAGTG | |
| *ndhG*-F | ATGGATTTACCTGGGCCAAT | |
| *ndhG*-R | TTATTGCCGAGCCATAGTAA | |
| *rpl2*-F | ACGGCGAAACATTTATACAA | |
| *rpl2*-R | TTACTTACGGCGACGAAGAATA | |
| *ropB-F* | ACTAAGCGTGCTATTCTCAA | |
| *ropB-R* | TTTATGGTCTAATTCCGAGC | |
| *rps8-F* | ATGGGCAAGGACACTATTG | |
| *rps8-R* | AACATAAGACTTCTCCCCCA | |
| *rps14-F* | ATGGCAAAAAAAAGTTTGATTC | |
| *rps14-R* | TTACCAACTGGATCTTGTTGCA | |
| *ycf3-F* | ATGCCTAGATCCCGTATAAATG | |
| *ycf3-R* | TTATTCAAATTCAAAGCGCTTC | |
